# Axial skeleton anterior-posterior patterning is regulated through feedback regulation between Meis transcription factors and retinoic acid

**DOI:** 10.1101/2020.03.09.983106

**Authors:** Alejandra C. López-Delgado, Irene Delgado, Vanessa Cadenas, Fátima Sánchez-Cabo, Miguel Torres

## Abstract

Vertebrate axial skeletal patterning is controlled by coordinated collinear expression of *Hox* genes and axial level-dependent activity of Hox protein combinations. Transcription factors of the Meis family act as cofactors of Hox proteins and profusely bind to Hox complex DNA, however their roles in mammalian axial patterning have not been established. Similarly, retinoic acid (RA) is known to regulate axial skeletal element identity through the transcriptional activity of its receptors, however, whether this role is related to Meis/Hox activity in axial patterning remains unknown. Here we study the role of Meis in axial skeleton formation and its relationship to the RA pathway by characterizing *Meis1*, *Meis2* and *Raldh2* mutant mice. Meis elimination produces axial skeleton defects without affecting Hox gene transcription, including vertebral homeotic transformations and rib mis-patterning associated to defects in the hypaxial myotome. While Raldh2 and Meis positively regulate each other, *Raldh2* elimination largely recapitulates the defects associated to Meis-deficiency and Meis overexpression rescues the axial skeletal defects in *Raldh2* mutants. We propose a Meis-RA positive feedback loop whose output is Meis levels and is essential to establish anterior-posterior identities and pattern of the vertebrate axial skeleton.

## INTRODUCTION

Anterior-posterior (AP) patterning is an essential feature of the bilaterian body plan and its mechanisms have been extensively studied. Canonical examples of AP patterning in vertebrates are found in the hindbrain and in the axial musculoskeletal system (Krumlauf, 1994). Segmental epithelial sacs known as somites emerge from the paraxial mesoderm as it is produced and progressively incorporate to the AP axis. The initially homogeneous somites later subdivide in compartments, including the sclerotome, precursor of the vertebrae and ribs, and the myotome, precursor of the skeletal muscles (Musumeci et al., 2015). Crosstalk from the myotome to the sclerotome is essential for sclerotome patterning and in particular for rib specification and patterning (Vinagre et al., 2010).

An important breakthrough in understanding antero-posterior axis patterning was the identification of *Hox* mutants in *Drosophila*, which cause the transformation of one part of the body into another, a phenomenon known as homeotic transformation (Lewis, 1978). *Hox* genes are conserved in evolution and appear organized in genetic complexes in most animals (Duboule and Dolle, 1989; Sanchez-Herrero et al., 1985). Mammals show *Hox* genes organized in 4 paralogous complexes (*HoxA*, *B*, *C* and *D*) that originated from two consecutive rounds of genome duplication and contain up to 13 paralogous genes. The genomic organization of Hox complexes correlates with their temporal and spatial expression domains, a phenomenon known as collinearity (Gaunt et al., 1986; Lewis, 1978). Mutations in *Hox* genes in different species produce AP homeotic transformations, which in mammals is best exemplified in the hindbrain and in the axial skeleton (Krumlauf, 1994).

*Hox* gene transcription is activated sequentially in axial precursors during gastrulation (13). Expression of *Hox* genes located at the 3’-most region of the complexes starts in axial progenitors in the posterior epiblast and is maintained in their descendants as they gastrulate through the primitive streak and colonize the embryonic AP axis. 3’-to-5’ sequential transcriptional activation of *Hox* complexes progresses continuously in axial progenitors, whereas their daughter cells fix their *Hox* expression code as they exit the progenitor region and colonize the embryonic axis. As cells colonize the different AP segments, they carry the successive Hox expression combinations to the progressively forming body axis, resulting in an AP nested patterns (Alexander et al., 2009). Thus, temporal information is translated into spatial domains during axial elongation (Deschamps and Duboule, 2017).

Hox proteins bind DNA through a 60 amino acid region called the homeodomain (McGinnis et al., 1984). The homeodomain is highly conserved and diversified in several transcription factor families in animals and plants. Hox proteins alone show limited DNA-binding ability, but they gain specificity and affinity for target sequences through interactions with cofactors of the PBC and MEINOX families, both belonging to the three amino acid loop extension (TALE) class of homeodomains (Mann and Affolter, 1998). PBC and MEINOX proteins form heterodimers and heterotrimers with Hox proteins, conferring them with increased target sequence selectivity and affinity (Merabet and Mann, 2016). In fact, mutants for the single members of the PBC and MEINOX families in *Drosophila* show AP phenotypes compatible with a generalized *Hox* loss of function, without affecting *Hox* AP expression (Chan et al., 1994; Rieckhof et al., 1997). In mammals, redundancy of the PBC (4 members) and MEINOX (5 members) families has hampered the study of their roles in axial skeletal patterning. While knowledge has been obtained from *Pbx* mutants in zebrafish and mouse, indicating essential roles in axial skeleton patterning (Capellini et al., 2008; Popperl et al., 2000; Selleri et al., 2001), the role of *Meis* genes in this context remains unexplored.

Meis proteins directly bind Hox proteins encoded by paralogs 9-13 (Shen et al., 1997) and form DNA-bound heterotrimeric complexes with Pbx and Hox proteins encoded by paralogs 1-10 (Chang et al., 1996). The repertoire of Meis, Prep and Pbx binding sites by ChIP-seq analysis in E11.5 mouse embryos identified Hox and Hox-PBC binding sites as the preferred sites for Meis binding, above the Meis-only binding sites, suggesting that Meis factors are strongly dedicated to interactions with Hox and Pbx proteins (Penkov et al., 2013). In addition, a large number of Meis binding sites was found within the *Hox* complexes, which suggested that additionally to their Hox-cofactor role, they may regulate *Hox* transcription. Studies in zebrafish (Choe et al., 2014) and mouse (Amin et al., 2015) embryos indeed showed that some of these binding sites represent *Hox* auto-regulatory elements active in the neural tube and Hox transcription regulation by Meis factors has been demonstrated in neural tube patterning (Dibner et al., 2001; Vlachakis et al., 2001; Waskiewicz et al., 2001). Meis elimination or overexpression also affects Hox gene expression during limb skeletal patterning (Delgado et al., 2020; Mercader et al., 1999; Mercader et al., 2009; Rosello-Diez et al., 2014), however, this aspect has not been studied in axial skeleton patterning.

Another interesting pathway that connects Meis, Hox and axial patterning is that of vitamin-A. The active form of Vitamin-A, retinoic acid (RA), regulates gene expression during embryonic development by binding to nuclear receptors RARα, RARβ or RARγ (Rhinn and Dolle, 2012). *Meis* genes have been identified in screens for RA targets (Berenguer et al., 2020; Oulad-Abdelghani et al., 1997) and respond to RA fluctuations *in vivo* (Mercader et al., 2000). RA excess produces axial skeleton alterations and modifies the *Hox* AP expression domains (Kessel and Gruss, 1991) and mutations in RA-receptor genes result in homeotic transformations, however, the mechanism by which this takes place is not clear. While RAR binding sites have been described in *Hox* complexes (Marshall et al., 1996), and RA administration *in vitro* regulates *Hox* gene transcription (Deschamps et al., 1987), RA administration *in vivo* can lead to axial skeleton homeotic transformations without changes in *Hox* expression (Kessel, 1992) and changes in *Hox* expression in *Rar*-deficient mice have not been reported.

Here we study the role of Meis factors in axial skeleton formation and its relationship to the RA pathway by characterizing mouse genetic models of *Meis1*, *Meis2* and *Raldh2*. We dissect the regulatory and functional relationships between *Meis*, *Hox* and *Raldh2* and formulate a new model that explains the ability of RA to produce homeotic transformations without modifying *Hox* expression.

## RESULTS

### *Meis* gene expression during anterior-posterior axial patterning of the mouse embryo

We studied the mRNA expression pattern of *Meis1* and *Meis2*, the two *Meis* genes extensively expressed in paraxial and lateral mesoderm (Figure 1). We detected the earliest expression of *Meis2* in early-streak stage embryos in a posterior region of the embryo close to the boundary with the extraembryonic region (Figure 1H). This expression extends distally and anteriorly as development progresses (Figure 1I) and at the early-headfold stage, an anterior stripe of *Meis2* transcripts was found bilaterally close to the extraembryonic region, and continuous with its posterior expression (Figure 1J). At late-headfold stage, *Meis2* started to disappear from the posterior region (Figure 1K) and at E8, the posterior embryonic bud was devoid of *Meis2* transcripts (Figure 1L). *Meis1* expression started slightly later than *Meis2*, being first detected at the late-streak stage, bilaterally in the mesoderm close to the extraembryonic region (Figure 1B) and at the early-headfold stage, forming a stripe of expression, similar to *Meis2* anterior stripe at this stage (Figure 1C). Both *Meis1* and *Meis2* expression domains extend posteriorly into the lateral plate mesoderm at the late-headfold stage (Figure 1D and 1K), but high levels of *Meis1* transcripts were never observed in the posterior embryonic bud. Finally, at E8 *Meis1* and *Meis2* expression patterns converge to a similar expression pattern, being strongly expressed in paraxial and lateral plate mesoderm up to the pharyngeal region (Figure 1E and 1L). At this stage, expression of both genes is excluded from the posterior embryonic bud, whereas it appears in the presomitic mesoderm and adjacent regions precursor to the lateral plate mesoderm. This expression pattern is maintained at later stages, indicating that as new precursors from the posterior bud incorporate to the presomitic area, they activate *Meis1* and *Meis2* and this activity persists as they differentiate into paraxial and lateral plate mesoderm. To determine the early activation pattern of *Meis1* and *Meis2* in the embryonic germ layers, we studied Meis mRNA and protein distribution in sections (Figure 1). Detection of Meis proteins in sections with an antibody that recognizes the majority of embryonic isoforms but does not discriminate between Meis1 and Meis2 shows early expression in all three germ layers at early allantoic bud stage (Figure 1 G and 1N). Sections of the RNA *in situ* hybridization of both genes, showed that *Meis1* expression was not detected in the epiblast/ectoderm (Figure 1F), while *Meis2* expression affected the three germ layers (Figure 1M). This result suggests *Meis2* is activated in epiblast cells and its expression persists as they gastrulate to contribute to mesoderm. The early *Meis2* expression pattern thus resembles the activation pattern of *Hox* genes.

**Figure 1.**
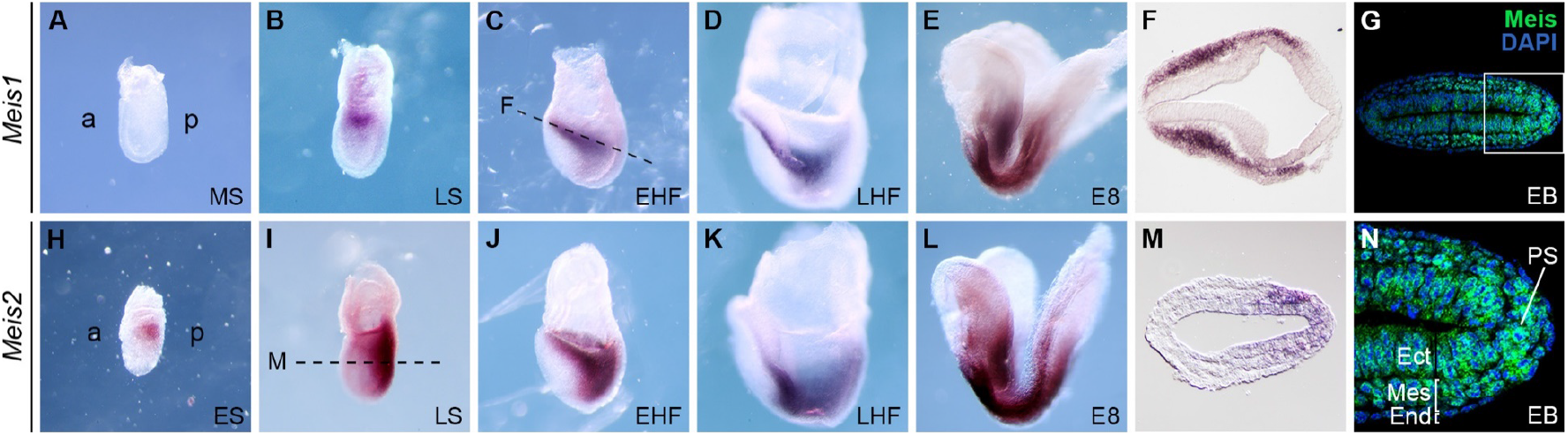
*Meis* expression pattern in early mouse embryo development. Whole-mount mRNA *in situ* hybridization of *Meis1* (A-E) and *Meis2* (H-L) from E7 to E8. (F and M) Transverse sections showing *in situ* hybridization for Meis1 and Meis2 mRNA, respectively. The approximate plane of the section is indicated by dashed lines in C and I. (G) Immunostaining with an antibody that recognized both Meis1 and Meis2 on longitudinal sections of an EB embryo across the PS. (N) Magnification of the region marked in G with the three germ layers indicated. a, anterior; p, posterior; MS, mid-streak; LS, late-streak; EHF, early headfold; LHF, late headfold; ES, early-streak; EB, early allantoic bud; PS, primitive streak; Ect, ectoderm; Mes, mesoderm; End, endoderm. All images are oriented with the anterior to the left and posterior to the right.

### *Meis* loss of function produces axial skeletal defects, including antero-posterior homeotic transformations

We used conditional deletion of *Meis1* and *Meis2* and studied the mutant skeletal pattern. We first studied the consequences of *Meis2* deletion using different Cre alleles that allow dissecting the putative specific functions of *Meis2* early expression. Deletion of a *Meis2^flox^* allele with *Sox2^Cre^* leads to *Meis2* elimination in the epiblast. Lethality of *Sox2^Cre^;Meis2^flox/flox^* embryos around E14.5-E15.5 due to cardiac defects did not allow us to study the pattern at later stages; however, the general vertebral formula could be determined at E14.5. We observed defects at the occipital-cervical transition, where the first cervical vertebra (C1 or atlas) was fused to the exoccipital bone (n=14/14) and in its ventral part showed a position and shape that resemble the exoccipital bone, while its dorsal part was not formed (Figure 2A and 2B; S1A Table). These changes correlated with a change in the shape of the second vertebra (C2 or axis), which acquired a C1-like morphology (n=13/14) (Figure 2A and 2B). With low penetrance, the C3 vertebra presented a morphology that resembles C2 (n=2/14). These observations are compatible with an anterior homeotic transformation of the cervical vertebrae. In addition, disconnected isolated elements often appeared (arrowhead in Figure 2B), suggesting as well segmentation problems in this region. Outside the axial skeleton, we observed a vestigial otic capsule in the mutants.

**Figure 2.**
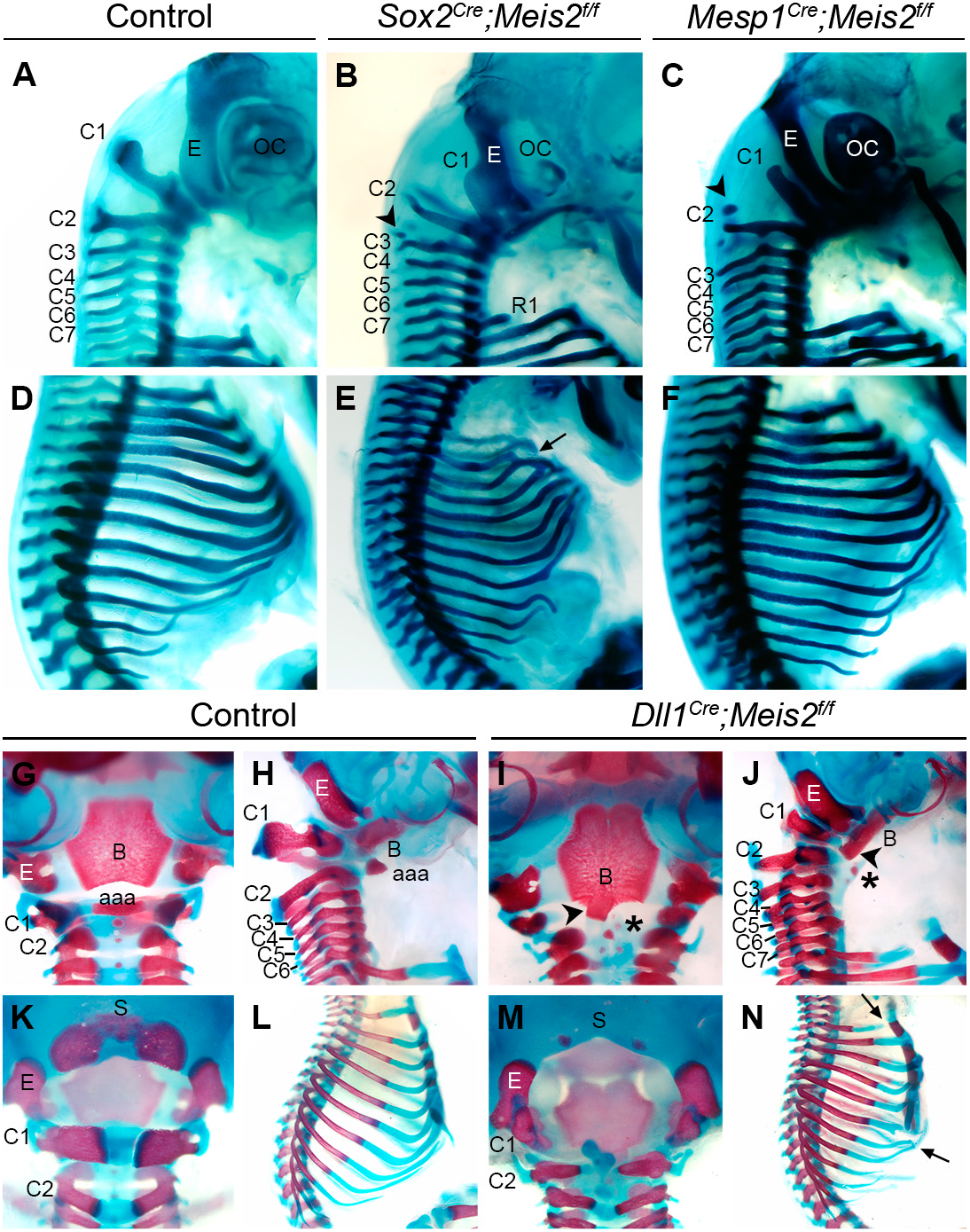
Skeletal defects in conditional *Meis2* mutant fetuses using different Cre alleles. (A-F) Victoria blue-stained skeletal preparations of E14.5 fetuses. The cervical region is shown for control (A), *Sox2^Cre^;Meis2^f/f^* (B) and *Mesp1^Cre^;Meis2^f/f^* (C) fetuses. Arrowheads in B and C point to disconnected chondrogenic condensations. The thoracic region is shown for control (D), *Sox2^Cre^;Meis2^f/f^* (E) and *Mesp1^Cre^;Meis2^f/f^* (F) fetuses. Arrows in E and N point to rib defects. (G-N) alizarin red/alcian blue-stained skeletal preparations of E18 control (G, H; K and L) and *Dll1^Cre^;Meis2^f/f^* (I, J, M and N) fetuses. The occipital region is shown in ventral (G and I), lateral (H and J) and dorsal (K and M) views. (L and N) lateral views of the thoracic region. Arrowheads in I and J indicate fusion between the basioccipital and the aaa. Asterisks in I and J indicate ectopic aaa formed on C2. aaa, anterior arch of the atlas; B, basioccipital; C, cervical vertebra; E, exoccipital; OC, otic capsule; R, rib; S, supraoccipital.

In the thoracic region, the most prominent defect was rib, rib-sternum attachment and sternum mispatterning (Figure 2D and E). We observed failures in sternum fusion, rib bifurcations, fusions and alteration of the sternal/floating rib formula. The gain of a rib in the first lumbar vertebra (L1) in some specimens (n=4/14) and the tendency to reduction of the first rib (R1) suggests the anterior transformations observed in the cervical region may also affect the thoracic region (Figure 2B). More caudal regions did not show any defects.

To investigate if *Meis2* activity in the epiblast is involved in the observed defects, we combined the *Meis2^flox^* allele with *Mesp1^Cre^* to eliminate *Meis2* from the nascent mesoderm. While *Mesp1* activates in the early embryo in a similar pattern to *Meis2*, because of the time lag between Cre expression and effective recombination, the recombination pattern of *Mesp1^Cre^* affects only the mesoderm and it does so down to the forelimb level (Saga et al., 1999). As it occurred with *Sox2^Cre^;Meis2^flox/flox^* mice, lethality due to cardiac defects only allowed us to study the phenotype at E14.5. In the *Mesp1^Cre^* model, we observed lower penetrance, but the same type and distribution of defects found in the *Sox2^Cre^* model, excepting the reduction of R1 and the otic capsule defects (Figure 2C and 2F; S1A Table).

To further dissect the specific tissues in which *Meis2* activity is required during early embryogenesis, we studied a third model in which we deleted *Meis2^flox^* using *Dll1^Cre^,* a line that recombines the mesoderm in the presomitic region (Wehn et al., 2009), i.e., at a later step of mesodermal allocation than *Mesp1^Cre^* does. *Dll1^Cre^;Meis2^flox/flox^* mice survive to adulthood, allowing a full assessment of the skeletal pattern at the end of gestation. In this model, we observed similar defects as those previously observed in the *Sox2^Cre^* and *Mesp1^Cre^* models in the occipital, cervical and thoracic regions (Figure 2G-N and 3M; S1A Table). In addition, we observed a defect in supraoccipital ossification (Figure 2K, 2M and 3M) and fusions between the basioccipital and the anterior arch of the atlas (aaa) (Figure 2G, 2I and 3M), which could not be determined at earlier stages because these bones form late in gestation. Again, we did not detect the formation of a rib in L1, suggesting this phenotype requires an early deletion of *Meis2*.

The irrelevance of early *Meis2* expression for most aspects of axial patterning is not due to compensatory activation of *Meis1*, as we detected no ectopic *Meis1* mRNA expression in early *Sox2^Cre^;Meis2^flox/flox^* embryos (S1 Figure). These results indicate that the expression of *Meis2* in the epiblast and early nascent mesoderm is to a large extent dispensable for its functions in axial skeletal patterning, although it might be needed for a proper specification of the thoracic-to-lumbar transition.

Next, to determine whether *Meis1* and *Meis2* cooperate in axial patterning, we combined *Meis1* and *Meis2* mutant alleles. Combining *Meis1* and *Meis2* deletion is not possible using the *Sox2^Cre^* or the *Mesp1^Cre^* deleters, due to lethality of double heterozygous mice. We therefore used the *Dll1^Cre^* line for these experiments. The defects observed in the allelic series generated affected the same skeletal elements that were altered in the *Meis2* mutant models (Figure 3) and the type of defects were similar, with anterior transformations of C1-C3 (Figure 3G-I and 3M; S1B Table) and defects in the occipital bones that either did not form or appeared fused to C1 or C2 element (Figure 3A-I and 3M; S1B Table). In the thoracic region we also detected rib fusions and defects in rib-sternum attachment (Figure 3J-L and 3M; S1B Table). Although we found 2 cases of extra ribs on L1, one case was also found in controls, suggesting this observation was unspecific. In general, skeletal defects are more severe as the number of *Meis* alleles deleted increases, being the absence of *Meis2* more detrimental than *Meis1* (Figure 3M; S1B Table). However, for some aspects of the phenotype, E18.5 *Dll1^Cre^;Meis1^flox/flox^;Meis2^flox/flox^* specimens appeared less affected in comparison with *Dll1^Cre^;Meis1^+/flox^;Meis2^flox/flox^* ones, which was paradoxical. We observed, however, that the viability of *Dll1^Cre^;Meis1^flox/flox^;Meis2^flox/flox^* mice at E18.5 was 37%, which suggested that specimens of this genotype at E18.5 represent escapers and thus, missing specimens could be more affected than appreciated. We then studied the phenotype of *Dll1^Cre^;Meis1^flox/flox^;Meis2^flox/flox^* fetuses at E14.5, when viability of double mutants was 67%, and observed a fraction of embryos with defects compatible with those observed at E18.5 and, in addition, we found very strongly affected fetuses showing all the cervical vertebrae fused, no apparent development of occipital condensations and widespread rib fusions and truncations (Figure 3N and 3O). In addition to the axial skeleton, defects in the limb skeleton were obvious in *Dll1^Cre^;Meis1^flox/flox^;Meis2 ^flox/flox^* fetuses (Figure 3N and 3O) and have been independently reported (Delgado et al., 2020).

**Figure 3.**
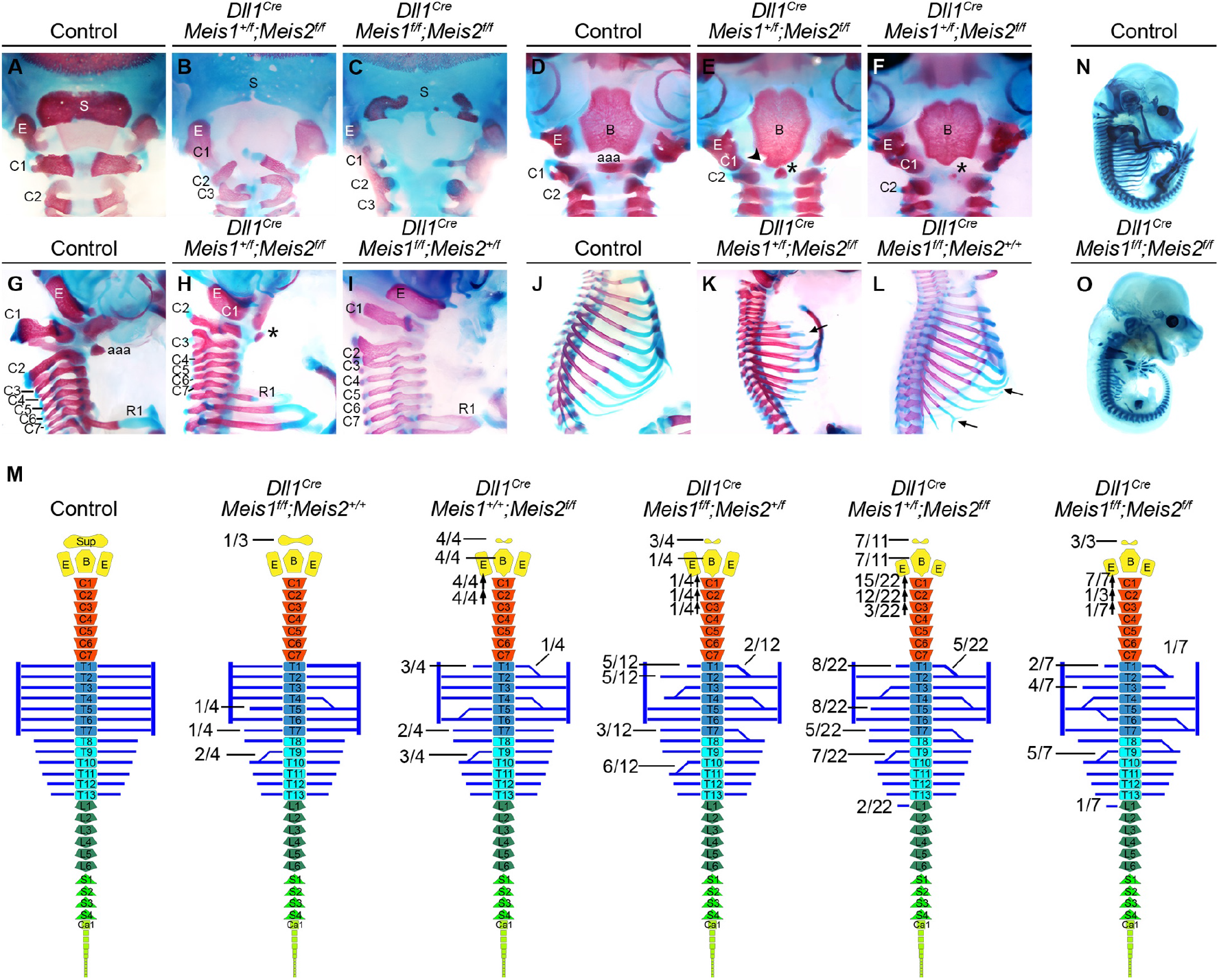
Skeletal defects in *Meis1* and *Meis2* loss of function mice using *Dll1^Cre^*. (A-L) alizarin red/alcian blue-stained skeletal preparations of E18.5 fetuses, control or mutant for different combinations of *Meis1* and *Meis2* floxed alleles and *Dll1^Cre^*, as indicated. The occipital region of control and mutant combinations is shown in dorsal (A-C), ventral (D-F) and lateral (G-I) views. (J-L) lateral views of the thoracic region. The arrowhead in E indicates a fusion between the basioccipital and aaa. Asterisks indicate ectopic aaa formed on C2. Arrows in K and L indicate rib defects. (M) Schematic representation of the axial skeleton defects of the different genotypes analyzed and their frequencies. Arrows pointing up indicate apparent anterior homeotic transformations. (N and O) Victoria blue-stained skeletal preparations of control and *Dll1^Cre^*-recombined *Meis1/2* homozygous floxed E14.5 fetuses. aaa, anterior arch of the atlas; B, basioccipital; C, cervical vertebra; E, exoccipital; R, rib; S, supraoccipital.

### *Hox* mRNA axial expression in *Meis* mutants

While defects in vertebral segmentation or overt rib deformations have not been described in Hox mutants, the AP specification defects observed in the occipital and cervical regions are very similar to those observed in the mutants of Hox paralog groups 3-5, and some of the rib cage defects are similar to those found in paralog groups 5-9 (Horan et al., 1995; Jeannotte et al., 1993; Manley and Capecchi, 1997; McIntyre et al., 2007). These coincidences and the previous reports of Meis proteins binding to *Hox* clusters (Amin et al., 2015; Penkov et al., 2013) prompted us to study the *Hox* mRNA expression pattern in *Meis* mutants. We did not detect alterations of *Hox* expression initiation or definitive anterior expression borders in either *Sox2^Cre^;Meis2^flox/flox^* or *Dll1^Cre^;Meis1^flox/flox^;Meis2^flox/flox^* embryos (Figure 4A and 4B). These results indicate that eliminating *Meis2* function with *Sox2^Cre^* or *Meis1* and *Meis2 Dll1^Cre^* does not modify *Hox* expression patterns and therefore, the phenotypes observed in these models do not relate to a Meis role in regulating *Hox* transcription. To study the generality of these observations, we combined simultaneous maternal and paternal deletion of *Meis1^flox/flox^* and *Meis2^flox/flox^* alleles, using the maternal deleter *Zp3^Cre^* and the paternal deleter *Stra8^Cre^* (S2 Figure). With this approach, we were able to completely eliminate *Meis1* and *Meis2* zygotic expression. Such embryos die around E9 with profound alterations of cardiac development, however this allowed us to study early *Hox* expression patterns. While previous reports in embryos at E9.5 or later stages have described paralog group Hox3 gene expression starting at somite 5 (Alexander et al., 2009), we found that in control embryos of up to 10 somites, expression of the Hox3 paralog group extends from somite 2-3 into more posterior somites (Figure 4C; S3 Figure). In embryos of 12 somites, the most anterior expressing somite is somite 3-4, while in embryos of more than 15 or more somites, expression starts at somite 5. These observations show transient Hox3 expression in occipital somites and a later progressive posteriorization towards their definitive expression domain. This expression pattern agrees with the fact that mutants of the group-3 Hox genes strongly affect the occipital region, which is mostly originated from somites 1-4. The defects present in group-3 Hox mutants in fact strongly affect the supraoccipital bone, which is exclusively contributed by somites 1 and 2 (Huang et al., 2001; Muller and O’Rahilly, 1994). Mutant embryos showed a normal Hox3-group gene expression in the paraxial mesoderm of embryos of 4-10 somites (Figure 4C; S3 Figure). Although counting somites was very difficult in mutant embryos of 15-20 somites, due to the developmental abnormalities, we concluded that the expression patterns in the paraxial mesoderm were either normal or anteriorized by 1-2 somites (Figure 4C; S3 Figure). In contrast, the anterior border of expression in the neural tube appeared clearly posteriorized (Figure 4C; S3 Figure). The study of the expression of *hoxd4* showed similar results, with a transient early expression starting at somite 4 and later getting restricted to its definitive anterior border at somite 6. In mutants, *hoxd4* expression was similar at early stages and appeared anteriorized to somite 4-5 at later stages. A posteriorization of Hox mRNA expression in the neural tube was again evident (Figure 4C). Most likely, the failure observed in the mutants in relocating *Hox* expression to more posterior somites at late stages does not indicate a direct role of Meis in regulating *Hox* expression, but a general blockade in development of *Meis* DKO embryos beyond the somite-7 stage. In fact, *Meis* DKO embryos do not undergo turning, body wall folding or neural tube closure, morphologically resembling E8.5 embryos at E9. The fact that no alterations were observed upon deletion with *Dll1^Cre^* support this conclusion.

**Figure 4.**
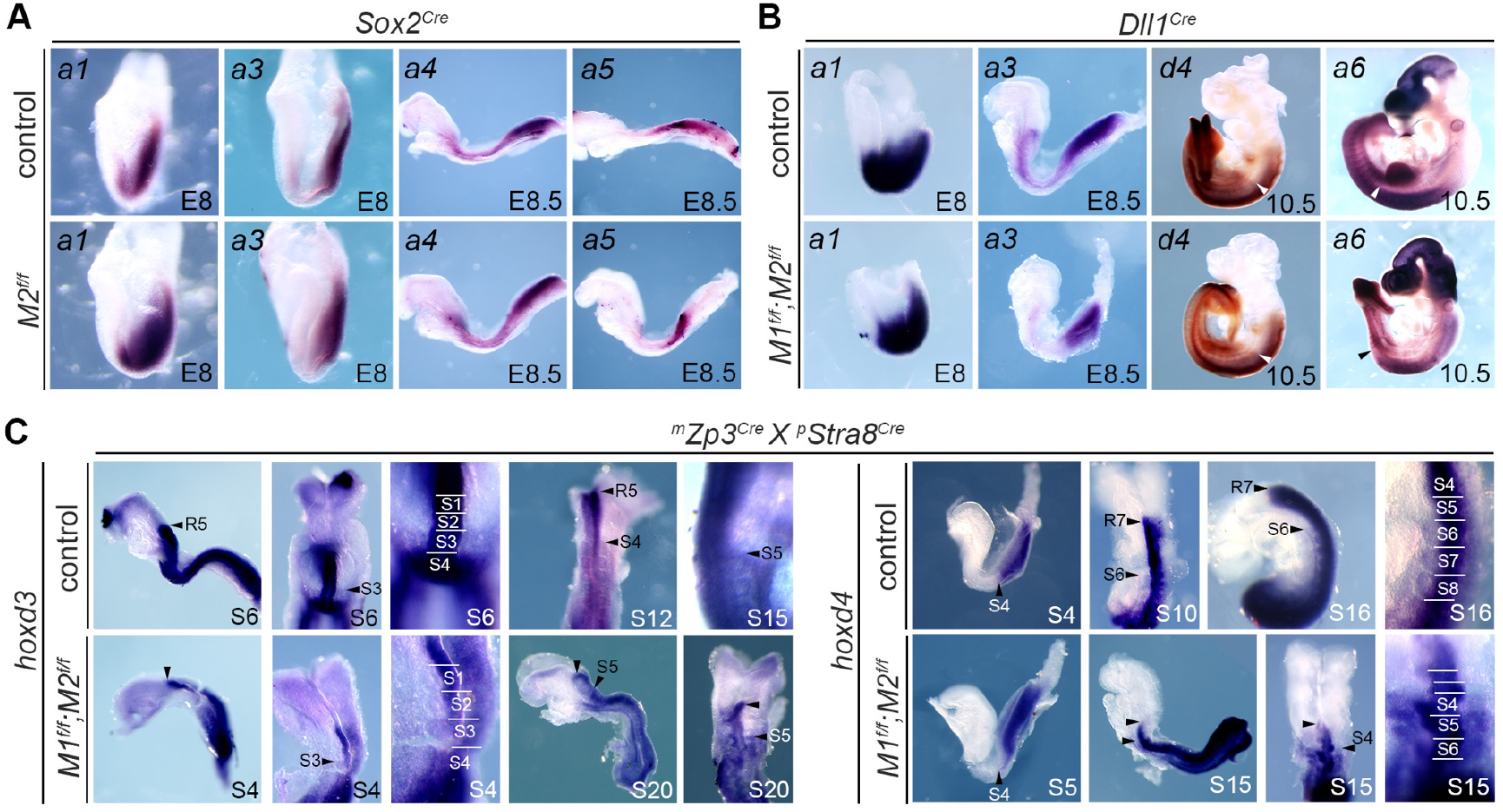
*Hox* gene mRNA expression patterns in *Meis* loss-of-function mutants. (A) mRNA *in situ* hybridization of the indicated *Hox* genes in E8-E8.5 control and *Sox2^Cre^*-recombined *Meis2* conditional mutant embryos. (B) mRNA *in situ* hybridization of the indicated *Hox* genes in E8-E10.5 control and *Dll1^Cre^*-recombined *Meis1* and *Meis2* conditional mutant embryos. (C) mRNA *in situ* hybridization of the indicated *Hox* genes in control and double-floxed *Meis1* and *Meis2* embryos derived from Zp3^Cre^ mothers and Stra8^Cre^ fathers.

We therefore conclude that transcriptional regulation of *Hox* genes is not involved in Meis regulation of axial skeleton patterning.

### Meis activity is required for hypaxial myotomal development

To identify the molecular mechanism underlying the skeletal phenotypes observed, we performed a transcriptomic analysis of E9 *Dll1^Cre^;Meis1^flox/flox^;Meis2^flox/flox^* embryos. To discriminate between alterations during the somite differentiation phase and early patterning defects, we separately analyzed the anterior region containing the first 10-12 somites and the posterior region including the rest of the somites and the posterior embryonic bud. We identified 9 upregulated genes and 25 downregulated genes in the analysis of the anterior region; whereas in the posterior region there were 58 upregulated and 58 downregulated genes (S4A and S4B Figure). Ingenuity Pathway Analysis showed that “skeletal and muscular system development” appears as the top tissue-specific altered class Figure S4C). Differences in other processes such as cell death, cell-to-cell interactions, cell assembly and organization were also found in this analysis (S4C Figure). No alterations were found in *Hox* gene expression, which confirmed the results observed in the *Hox* mRNA *in situ* analysis.

We then focused on the *in situ* analysis of genes involved in somite development found altered in the RNAseq analysis and in additional genes relevant to somite patterning. When comparing the expression pattern of this set of genes between control and *Dll1^Cre^;Meis1^flox/flox^;Meis2^flox/flox^* embryos, we found that a set of genes expressed and/or involved in hypaxial myotomal development were downregulated in the hypaxial region of the mutants (Figure 5), including *Eya1* (Grifone et al., 2007) (Figure 5A and 5B), *Sim1* (Ikeya and Takada, 1998) (Figure 5C and 5D), *Shisa2* (Nagano et al., 2006) (Figure 5E and 5F) and *Pax3* (Tremblay et al., 1998) (Figure 5G and 5H).

**Figure 5.**
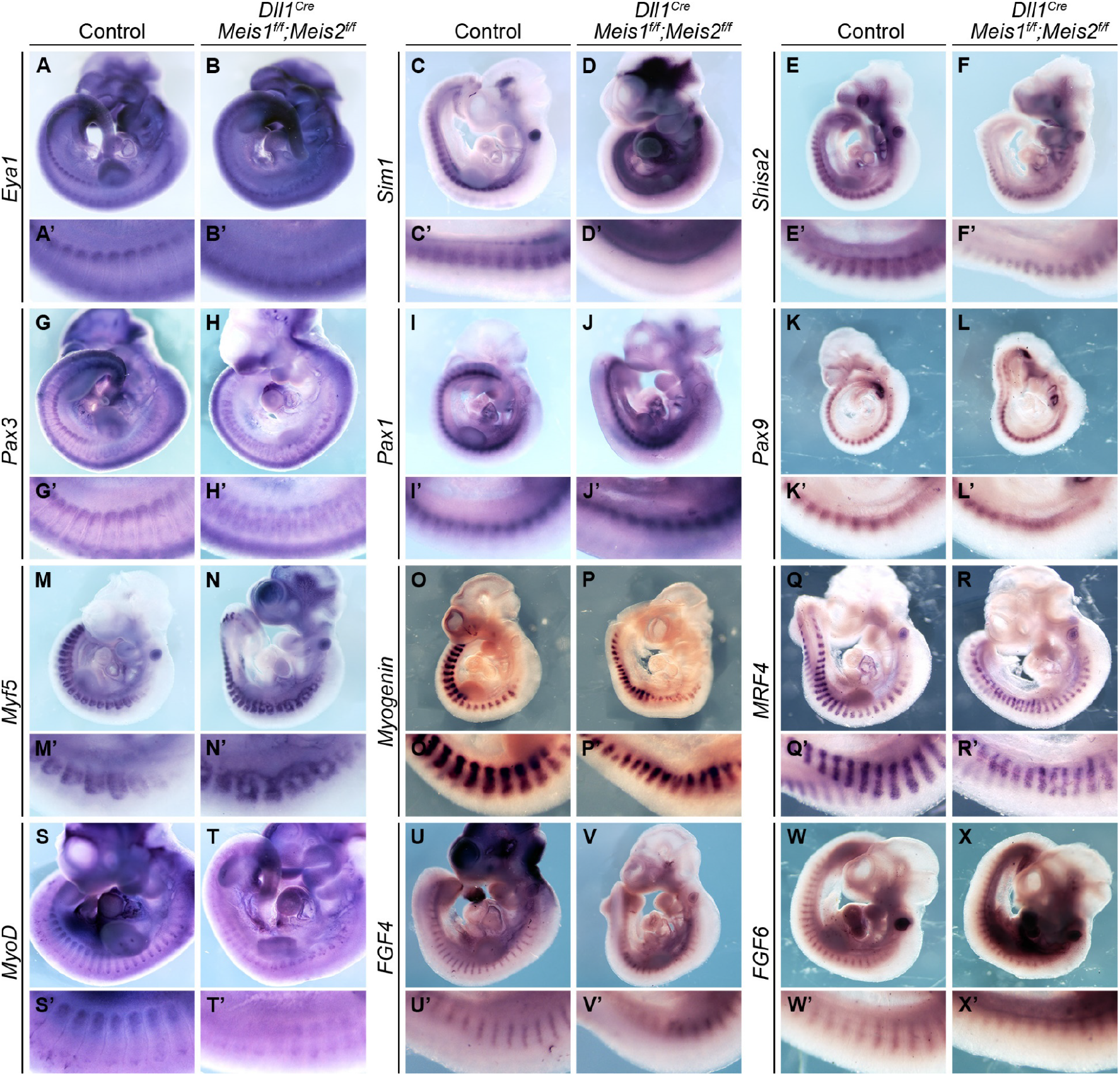
Expression analysis of genes involved in somite development in *Meis* mutants. Images show whole mount in situ mRNA hybridization in E10.5 embryos showing the expression of genes relevant for somitogenesis in control and *Dll1^Cre^;Meis1^f/f^;Meis2^f/f^* embryos, as indicated. (A and B) *Eya1*, (C and D) *Sim1*, (E and F) *Shisa2*, (G and H) *Pax3*, (I and J) *Pax1*, (K and L) *Pax9*, (M and N) *Myf5*, (O and P) *Myogenin*, (Q and R) *MRF4*, (S and T) *MyoD*, (U and V) *FGF4* and (W and X) *FGF6*. (A’-X’) Magnification of the trunk region of the corresponding image.

Regarding sclerotome markers, we found no alteration of *Pax1* expression (Figure 5I and 5J); however, an abnormal expression pattern of *Pax9* was observed in sclerotomes of the cervical region, which appeared incorrectly segmented (Figure 5K and 5L). Crosstalk between the myotome and sclerotome is essential for proper rib patterning and mice deficient for the myogenic factors Myf5 (Braun et al., 1992), MRF4(Zhang et al., 1995), or Myogenin (Vivian et al., 2000) show rib defects that resemble those described here in Meis mutants. Therefore, we next studied the main myogenic factors. Expression of *Myf5* appears first in the epaxial somite at E8, followed by *MRF4* and *Myogenin* at E9, later extending hypaxially caudal to the forelimb at E10.5 (Figure 5M-R). In mutant mice, the early epaxial expression of *Myf5* shows incomplete segmentation, whereas at E10.5, expression in myotomes anterior to the forelimb extends ventrally and appears as a continuous band between adjacent somites in a pattern that is not detected in control embryos (Figure 5M and 5N). Both *MRF4* and *Myogenin* show mis-segmented and bifurcating patterns in mutant embryos (Figure 5 O-R). In addition, the ventral hypaxial extension of the expression domain was reduced, as observed before for other hypaxial markers. In contrast, defects in the early expression of *Myogenin* at E9 are not as evident as for *Myf5*. *MyoD* shows as well a disorganized and spread expression in cervical myotomes of mutants, whereas hypaxial extension of the expression is also defective in more caudal myotomes (Figure 5T).

We finally studied *FGF4* and *FGF6*, which are involved myogenesis through their expression in the medial myotome (Grass et al., 1996). We found that expression of *FGF4* and *FGF6* appeared highly reduced in mutant embryos (Figure 5U-X).

In summary, re-segmentation of the paraxial mesoderm appears impaired in *Meis* mutants, with defects in the separation of adjacent sclerotomal/myotomal domains and bifurcated myogenic domains. These defects affect mainly the cervical region, although defects were also seen some times in the interlimb region. During further myotome development, a defect in myogenic *FGF* expression was found and the hypaxial developmental program seems especially affected with a failure in hypaxial myotomal migration, in correlation with an inability to properly activate *Pax3* expression.

### A positive feedback loop maintains the Retinoic Acid pathway and *Meis* expression during axial patterning

In the transcriptomic analysis of *Meis* mutants, *Raldh2* –the gene encoding the main enzyme responsible for embryonic RA synthesis–, and *Cyp26b1* –the gene encoding the main enzyme responsible for RA degradation in the embryo– appeared downregulated in the anterior trunk region by RNA-seq (S4 Figure). *In situ* hybridizations for both genes were consistent with the transcriptomic analysis. *Raldh2* expression appeared reduced at E9.5 in the differentiating derivatives of anterior somites but not in the presomitic area or in newly produced somites (Figure 6A-B’’). A similar pattern is present at E10.5, where tail regions with newly produced somites do not show alterations but more anterior regions do show a reduction in *Raldh2* transcripts (Figure 6C-D’’).

**Figure 6.**
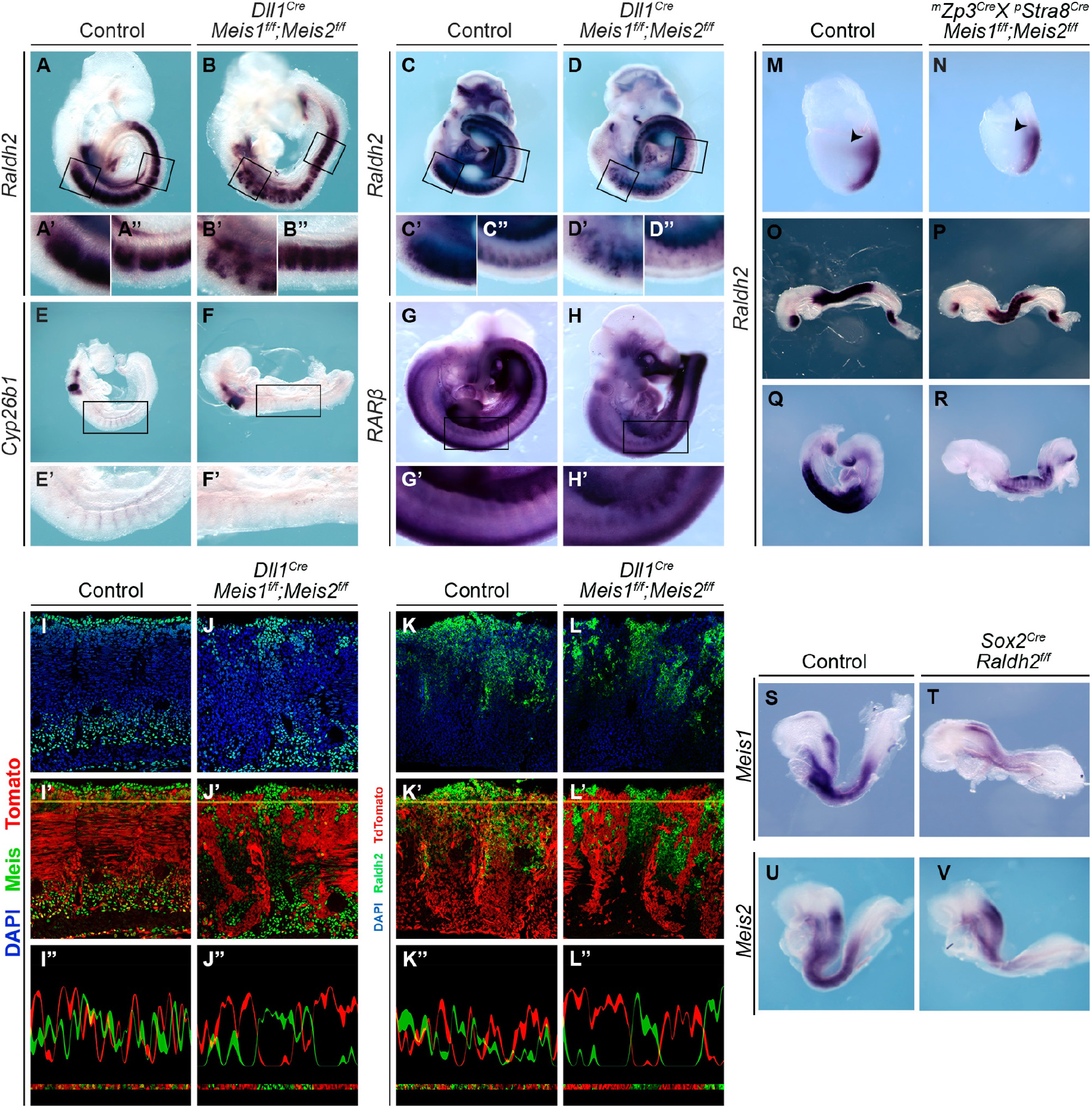
Cross-regulatory interactions between Meis and the Retinoic Acid pathway. (A-D’’) *Raldh2* mRNA *in situ* hybridization in E9.5 (A-B’’) and E10.5 (C-D’’) control and *Dll1^Cre^*-recombined double *Meis1* and *Meis2* mutant embryos, as indicated. E9.5. (E-F’) *Cyp26b1* mRNA *in situ* hybridization of control and *Dll1^Cre^;Meis1^f/f^;Meis2^f/f^* E9 embryos, as indicated. (G-H’) *RARβ* mRNA *in situ* hybridization of control and *Dll1^Cre^;Meis1^f/f^;Meis2^f/f^* E10.5 embryos, as indicated. (A’-H’ and A’’-D’’) Magnification of the regions marked in the upper images. (I-L’’) *Dll1^Cre^* recombination pattern reported by a *Rosa26R^tdTomato^* allele in the anterior somites of E10.5 embryos. Meis and Raldh2 immunofluorescence (I-L’) and corresponding quantification plots (I’’-L’’) along the indicated yellow lines in I’-L’, in control and *Dll1^Cre^;Meis1^f/f^;Meis2^f/f^* embryos, as indicated. (M-R) *Raldh2* mRNA *in situ* hybridization in embryos at E7.5 (M, N), E8.75 (O-P) and E9 (Q-R) of control (M, O and Q) and maternally and paternally-recombined *Meis1^f/f^;Meis2^f/f^* (N, P and R) embryos. Arrowheads indicate the *Raldh2* expression domain in the lateral plate in M and its absence in N. (S-V) *Meis1* and *Meis2* mRNA *in situ* hybridization in control (S, U) and *Sox2^Cre^;Raldh2^f/f^* (T, V) E8.5 embryos. *^m^Zp3^Cre^* indicates maternal presence of the allele and *^p^Stra8^Cre^* indicates paternal presence of the allele.

In the somitic region of E9 embryos, *Cyp26b1* is expressed exclusively in the endothelium of the dorsal aortae and inter-somitic vessels, whereas it is strongly expressed in areas of the hindbrain. The hindbrain signal was preserved in mutants; however, the endothelial signal in the trunk region was lost (Figure 6E-F’). *Cyp26b1* is a direct target of the RA pathway that gets activated in response to RA. The concomitant downregulation of *Raldh2* and *Cyp26b1* thus suggest that *Meis* mutant embryos are defective in RA. We then studied the expression of the gene encoding the RA receptor beta (*RARβ*), which has been described as a RA-responsive gene. Contrary to expectations, no change in the pattern of *RARβ* was detected between controls and *Meis* mutants (Figure 6G-H’), which is consistent with the RNAseq data that identified no differences in *RARα*, *RARβ* or *RARγ*.

Unexpectedly, several of the embryos studied showed *Raldh2* reduction in a mosaic fashion. To understand why the reductions in *Raldh2* appeared in a mosaic fashion, we combined *Meis1^flox^* and *Meis2^flox^* alleles with *Dll1^Cre^* and a *Rosa26R^tdTomato^* reporter. In these embryos, *Dll1^Cre^* recombines the *Rosa26R^tdTomato^* reporter, allowing to determine the Cre recombination pattern. At E10.5, E9.5 and E8.5, we observed a mosaic pattern of Tomato^+^ cell distribution in both control and *Meis* mutant embryos, with variability in the proportion of Tomato^+^ cells found in the somites of different embryos (Figure S5). This mosaicism had not been described for this line before (38) and therefore it might depend on the genetic background. To determine whether the observed mosaicism results from inefficient recombination in all cells or from mosaic activation of *Dll1^Cre^* expression, we studied the correlation between Meis immunodetection and Tomato expression in *Dll1^Cre^;Rosa26R^tdTomato^;Meis1^flox/flox^;Meis2^flox/flox^* embryos. We found that Tomato^+^ cells were devoid of Meis, while their neighboring Tomato^-^ cells showed Meis expression (Figure 6I-J’’). Image profiling shows anti-correlation between Tomato and Meis detection in mutants (Figure 6J’’), whereas this was not found in control embryos (Figure 6I’’). These observations indicate that the pattern observed results from mosaic inactivation of *Dll1^Cre^* and therefore the Tomato^+^ cell distribution reports the distribution of *Meis*-deficient cells. In mutants, we found a tendency of knockout and wild type cells to segregate from each other, resulting in large aggregates of Tomato^+^ cells that were not found in controls (Figure 6I’ and 6J’). We did not find any reproducible difference between mutant and control embryos in the distribution of Tomato^+^ cell patches by tissues. In addition, the anterior-most border of Tomato^+^ cell distribution was established at the occipital level and this was not different between control and mutant embryos.

We therefore used the mosaic inactivation of *Meis* alleles to study the regulation of *Raldh2* by Meis. We performed Raldh2 immunostaining and correlated this signal with that of Tomato. We found that Tomato^+^ cells lacking Meis function did not present detectable Raldh2 expression, while their Tomato^-^, Meis-expressing, neighboring cells showed normal Raldh2 expression (Figure 6K-L’’). The result was similar to that observed for Meis immunostaining, being the signal of Raldh2 and Tomato mutually exclusive in mutant embryos but not in controls (Figure 6K’’ and 6L’’). These results indicate a strict and cell-autonomous requirement of Meis function for *Raldh2* expression in the differentiating trunk mesoderm.

We then analyzed *Raldh2* expression in embryos with double maternal/zygotic inactivation of *Meis1* and *Meis2* (Figure 6M-R). *Raldh2* mRNA distribution in the early embryo resembles *Meis* expression pattern; however, it starts slightly later and only affects the mesoderm (Figure S6). In mutant embryos, we observed no alteration of the expression pattern in the axial and paraxial mesoderm, however the lateral plate domain close to the extraembryonic region was abolished (arrowheads in Figure 6M and 6N). Up to E8.75, when only the first somites have formed, no alteration of *Raldh2* expression in the paraxial mesoderm is observed (Figure 6O and 6P), however at E9 all trunk *Raldh2* expression is strongly decreased in mutants (Figure 6Q and 6R).

Given that retinoic acid has been shown to regulate *Meis* expression in different settings (Mercader et al., 2000; Oulad-Abdelghani et al., 1997; Yashiro et al., 2004), we studied whether the elimination *Raldh2*-mediated RA synthesis affects axial Meis expression. We studied *Meis1* and *Meis2* mRNA and protein expression in *Sox2^Cre^; Raldh2^flox/flox^* embryos and found that both genes presented a reduction of transcripts along the trunk region of E8 embryos (Figure 6S-V).

These results indicate that Meis is required for maintenance of *Raldh2* expression in the differentiating paraxial mesoderm but not for its initial expression before somite differentiation. These conclusions correlate with the observed downregulation of *Raldh2/Cyp26b1* in the transcriptome of anterior trunk but not the posterior trunk of E9.5 embryos. In contrast, the early lateral plate mesoderm –likely fated to the cardiogenic area– requires Meis activity for *Raldh2* expression from the earliest stages. Reciprocally, *Raldh2* expression is required to maintain proper *Meis* expression levels, but not for initiating *Meis* expression, given that *Meis* expression starts before *Raldh2* expression. These results indicate that there is a positive regulatory loop between *Meis* and *Raldh2* that is relevant to mutually maintain but not initiate their expression.

### *Raldh2* deficiency produces axial skeleton defects partially overlapping with those observed in *Meis* mutants

While retinoic acid has long been postulated as a regulator of axial skeleton, there is no direct study of the consequences of eliminating RA on antero-posterior axial identities. Here, we conditionally deleted *Raldh2* using *Dll1^Cre^* to investigate whether this affects the axial skeleton and the extent to which RA might be related to Meis roles in axial patterning. In the occipital region, the basioccipital presented similar alterations to those observed in *Meis* mutants (n=14/43) (Figure 7A and 7F; S1D Table), including its fusion with the aaa (Arrowhead in Figure 7F). Strikingly, similar modifications of the basioccipital were also found in some control embryos, although in a lower proportion (n=5/47) (Figure 7P), suggesting a genetic background prone to these particular defects. In mutants, C1 appeared fused to, and/or adopting a shape and position similar to the exoccipital (N=8/49). In the cases in which C1 showed transformation to exoccipital, C2 adopted a C1 morphology (n=8/41), whereas some cases in which C1 retained its morphology, C2 adopted a C1 morphology and partially fused to C1 (n=9/41). C3 to C2 transformations/fusions were also observed (n=11/41). At the cervical thoracic transition, tuberculi anterior were found in C7 instead of C6 (n=5/21) (Figure 7C and 7H, arrow in 7H), suggesting that anterior transformations also take place at this axial level.

**Figure 7.**
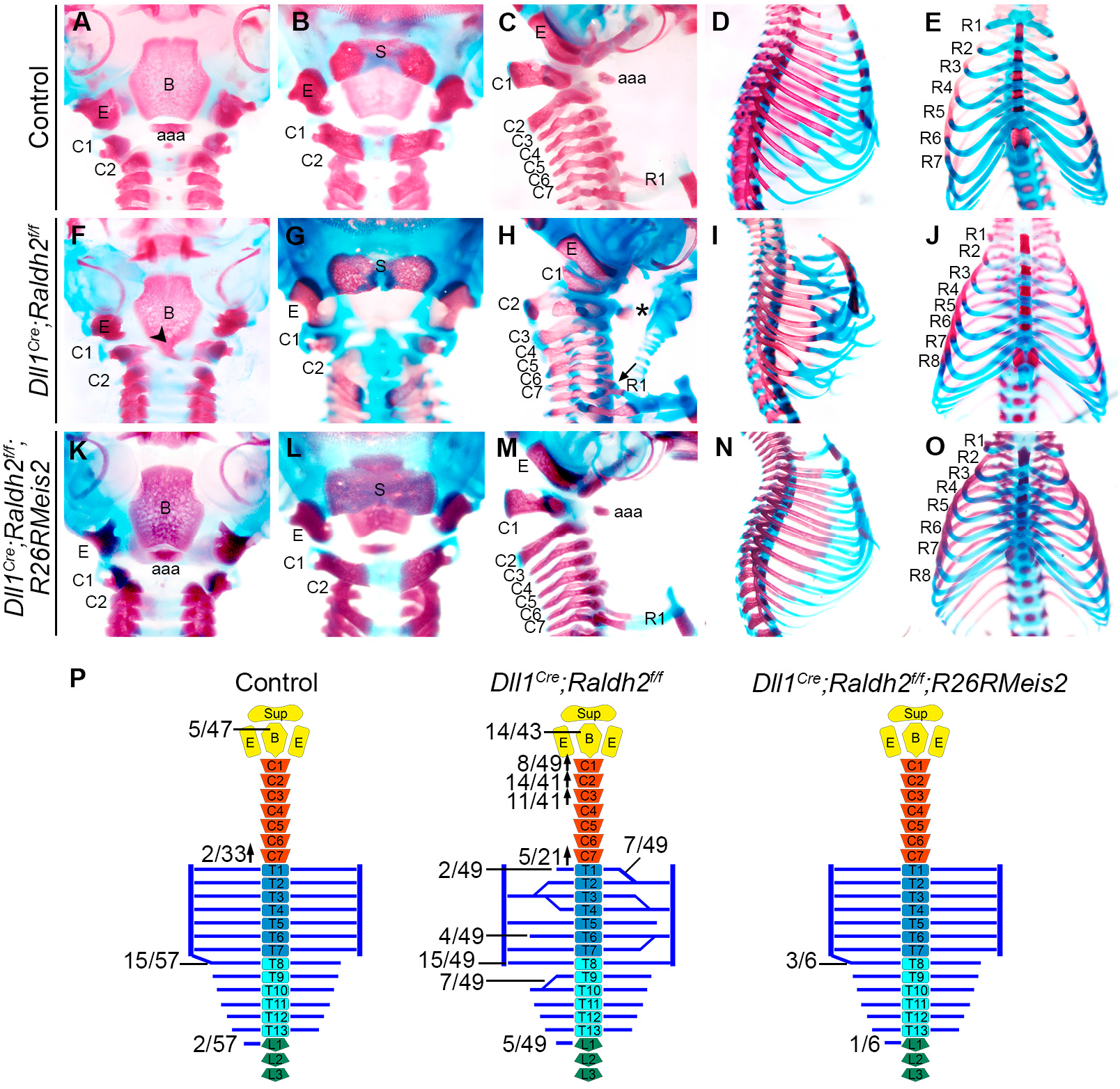
Skeletal defects in *Dll1^Cre^;Raldh2^f/f^* fetuses and their rescue by Meis2 expression. (A-J) Skeletal staining of E18.5 fetuses. Ventral view of the basioccipital in control (A) and *Dll1^Cre^;Raldh2^f/f^* fetuses (F). Arrowhead in F points fusion between basioccipital and aaa. Dorsal view of the supraoccipital in control (B) and *Dll1^Cre^;Raldh2^f/f^* fetuses (G). Cervical region in control (C) and *Dll1^Cre^;Raldh2^f/f^* fetuses (H). Asterisk in H indicate aaa formed by C2 and arrow point to tuberculi anterior in C7. Thoracic region in control (D) and *Dll1^Cre^;Raldh2^f/f^* fetuses (I). Ventral view of the sternum in control (E) and *Dll1^Cre^;Raldh2^f/f^* fetuses (J). (K) Schematic representation of the axial skeletal defects of *Dll1^Cre^;Raldh2^f/f^* fetuses and their frequencies. Upward and downward arrows respectively indicate anterior or posterior homeotic transformations. aaa, anterior arch of the atlas; B, basioccipital; C, cervical vertebra; E, exoccipital; R, rib; S, supraoccipital.

Altogether, the alterations found in *Raldh2* mutants in the occipital/cervical regions were similar to those observed in *Meis* mutants but displayed lower penetrance (Figure 7P). In the thoracic region, shortening or fusion of the first rib with the second rib and generalized rib fusions and bifurcations were observed in similarity to the defects found in Meis mutants (Figure 7D and 7I). In the most affected mutant embryos, we observed defects in the inter-sternal cartilage and the sternebrae, although we did not observe a split sternum (Figure 7E and 7J). Some incidence of an extra sternal rib and an extra rib on L1 was also observed (Figure 7E, 7J and 7P), suggesting A-P transformations were extensive down to the thoracic/lumbar transition.

The compared analysis of *Meis* and *Raldh2* mutants supports the idea that Meis and the retinoic acid pathway act in a positive feedback loop that is relevant in patterning the axial skeleton. To obtain evidence for the functional relevance of this regulatory loop and determine its relevant output in axial patterning, we used a *Rosa26R^Meis2^* allele that provides *Meis2* overexpression upon Cre recombination. We then simultaneously eliminated *Raldh2* and activated *Meis2* with *Dll1^Cre^*. Interestingly, in this mouse model, all defects produced by *Raldh2* mutation in the axial skeleton were rescued (Figure 7K-O, 7P; S1D and S1E Table), indicating that Meis suppresses the effect of RA deficiency on axial skeleton patterning.

## DISCUSSION

*Meis1* and *Meis2* expression starts at gastrulation, although their early patterns are different in time and expression domains. *Meis2* is activated earlier than *Meis1* in a pattern that coincides spatially and temporally with that of *Hox* gene activation in the posterior epiblast. Despite this, we have not observed alterations in *Hox* gene expression patterns or transcript abundance in *Meis* mutants. These results indicate that, despite the profuse binding of Meis proteins to the Hox complexes (Penkov et al., 2013), Meis is not involved in *Hox* gene transcriptional regulation during axial skeleton patterning. This is not extendable to other embryonic regions, given that we have observed clear alterations of *Hox* mRNA expression domains in limb buds with altered Meis function (Delgado et al., 2020; Mercader et al., 1999; Mercader et al., 2009; Rosello-Diez et al., 2014) and similar results have been reported in neural patterning (Dibner et al., 2001; Vlachakis et al., 2001; Waskiewicz et al., 2001). The observed binding of Meis to the Hox complexes therefore might be involved in regulating Hox transcription in several tissues but not in the paraxial mesoderm, at least at the stages studied here.

Despite the absence of changes in *Hox* transcription in the paraxial mesoderm, *Meis* mutants produce anterior homeotic transformations and defects similar to those previously described for *Hox* mutants involved in patterning the occipital, cervical and thoracic regions (Horan et al., 1995; Jeannotte et al., 1993; Manley and Capecchi, 1997; McIntyre et al., 2007). This is consistent with studies in flies in which the elimination of the *Meis* ortholog *homothorax* produces homeotic phenotypes through modifying Hox protein DNA affinity and target selectivity without altering *Hox* gene transcription (Merabet and Mann, 2016; Rieckhof et al., 1997).

We deleted *Meis2* using different Cre lines that recombine at different stages of epiblast cell recruitment to the paraxial mesoderm, however we did not find any substantial influence of the timing of *Meis2* removal on the phenotypes obtained. Using *Dll1^Cre^*, which recombines in the pre-somitic mesoderm, the severity of the defects observed increases with the number of *Meis* alleles deleted, supporting a cooperation between *Meis1* and *Meis2* in axial patterning. The early expression of *Meis2* in the posterior epiblast thus seems not to play any role in axial skeletal patterning, while both *Meis1* and *Meis2* cooperate at the presomitic mesoderm, or at later stages of somite development, in axial patterning. Although it has been suggested that segmental identity specification occurs in the PSM before somites are formed (Carapuco et al., 2005), we cannot exclude the involvement of Meis during later somite development, given that Meis is also present in the differentiating paraxial mesoderm and that we have not eliminated Meis function specifically from the differentiating somites.

Apart from its function in segmental identity, the transcriptional analysis of the mutants indicates an important function in hypaxial myotome development, with profound alterations of both patterning and myogenic pathways. Interestingly, *Myf5*, *MRF4* and *Myogenin*-deficient mice show rib defects similar to those described here (Braun et al., 1992; Hasty et al., 1993; Patapoutian et al., 1995), and therefore, the failure in proper activation of the hypaxial myogenic program is sufficient to explain rib mispatterning in *Meis1/2* double KOs. Moreover, hypaxial myotomal *FGF4* and *FGF6* expression, required for rib patterning downstream the myogenic factors (Vinagre et al., 2010), is strongly impaired in *Meis* mutants, indicating a function of Meis in the cross-talk between myotome and sclerotome. In addition, the activation of the myogenic program involved in rib pattering is under direct control of a specific set of Hox proteins involved in the specification of thoracic segments (Vinagre et al., 2010). The rib mispatterning phenotypes therefore may also partly involve the impairment of Hox function in the absence of Meis.

RNA-seq analysis and *in situ* hybridization revealed a reduction in *Raldh2* and *Cyp26b1* in *Meis* mutants. Since the activation of *Cyp26b1* is RA-dependent, its downregulation in *Meis* mutants could be a secondary event, due to the reduction in RA synthesis by Raldh2. *Cyp26b1* mutants show posterior homeotic transformations in the occipital/cervical region (Sakai et al., 2001), associated to increased RA levels. The posterior transformations in this model are opposite to those observed in *Meis* mutants, which concurs with the idea that Meis is a positive regulator of RA synthesis. In addition, *in vivo* treatments with RA during mouse gestation caused either anterior or posterior homeotic transformations depending on the stage of the treatment (Kessel and Gruss, 1991). In the cervical region, anterior transformations were observed following treatments at E7 while posterior transformations were found following RA treatment from E8 (Kessel, 1992). On the other hand, *Raldh2* knockout mice die around E10.5 from an impairment in RA synthesis (Niederreither et al., 1999), however, a conditional approach that would allow studying the skeletal pattern was missing. We generated a *Raldh2* conditional knockout using the *Dll1^Cre^* driver and found homeotic transformations affecting the occipital/cervical region, and additional patterning defects in the thoracic region that significantly overlap with those observed in *Meis* mutants. In agreement with this, mutations in RARs lead to homeotic transformations (Lohnes et al., 1993; Lohnes et al., 1994) similar to those observed in *Meis* mutants. In particular, *RARγ* and *RARβ* loss of function mutants show anterior transformations without showing any changes in *Hox* expression patterns (Folberg et al., 1999a; Folberg et al., 1999b).

At the molecular level, we described a positive regulatory loop between Meis and RA that we confirmed by functional genetic analysis. The similarities in skeletal transformations between *Raldh2* and *Meis* mutants and the cross-regulation between *Raldh2* and *Meis* suggests that the positive regulatory loop between *Raldh2* and Meis is involved in axial patterning. While *Meis* genes are RA targets in various contexts (Mercader et al., 2000; Oulad-Abdelghani et al., 1997; Yashiro et al., 2004), *Raldh2* is a direct Meis target in the hindbrain (Vitobello et al., 2011), and ChIPseq analysis in E10.5 limb buds (Delgado et al., 2020) identifies Meis binding sites in the *Raldh2* locus. In fact, Meis could promote RA accumulation at various levels, as it also represses *Cyp26b1* during limb development in a cell-autonomous manner (Rosello-Diez et al., 2014). The requirement of Meis activity for *Raldh2* transcription in the paraxial mesoderm is restricted to the differentiation stages and does not take place in the nascent or segmenting mesoderm. In coincidence with our findings, *Pbx1/2* null embryos show normal *Raldh2* expression at early embryonic stages but strong downregulation in the paraxial mesoderm at E9.0 and beyond (Rosello-Diez et al., 2014).

Finally, we studied the functional output of the Meis-RA regulatory loop by genetic rescue. The complete rescue of *Raldh2* mutants by *Meis* overexpression suggest that Meis is the main functional output of the positive regulatory loop between Meis and RA in the paraxial mesoderm.

We propose a model for the RA-Meis-Hox network in the paraxial mesoderm in which Meis is involved in a positive feedback loop with RA through *Raldh2* regulation. Meis is the main output of this regulatory loop and is required for the specification of axial skeletal identities, likely through regulating Hox protein activity. The proposed model provides a possible explanation to the ability of RA and RARs to phenocopy Hox mutants without affecting their transcriptional expression.

## MATERIALS AND METHODS

### Mouse lines and embryo harvest

Experiments were performed using mice (*Mus musculus*). Mice were handled in accordance with CNIC Ethics Committee, Spanish laws and the EU Directive 2010/63/EU for the use of animals in research. All mouse experiments were approved by the CNIC and Universidad Autónoma de Madrid Committees for “Ética y Bienestar Animal” and the area of “Protección Animal” of the Community of Madrid with reference PROEX 220/15.

*Meis* conditional knockouts were generated mating *Meis1^flox^* (Unnisa et al., 2012) and *Meis2^flox^* (Delgado et al., 2020) with different Cre lines: *Sox2^Cre^* (Hayashi et al., 2002), *Mesp1^Cre^* (Saga et al., 1999), *Dll1^Cre^* (Wehn et al., 2009), *Stra8^Cre^* (Sadate-Ngatchou et al., 2008) and *Zp3^Cre^* (de Vries et al., 2000). *Raldh2* conditional knockouts were obtained by mating *Raldh2^flox^* mice (79) to *Dll1^Cre^* and *Sox2^Cre^*. For Cre^+^ cell lineage tracing we used *Gt(ROSA)^26Sortm14(CAG-tdTomato)Hze^* (Madisen et al., 2010). For conditional Meis overexpression we used the *Rosa26R^Meis2-EYFP^* line (Rosello-Diez et al., 2014).

To obtain embryos at different gestational stages, mice were mated in the afternoon and females were checked every morning for the presence of a vaginal plug; noon of the day the plug was observed and considered as gestational day 0.5 (E0.5). Embryos at somitogenic stages were staged according to age and somite number. Embryos that had not started somitogenesis were staged according to (Downs and Davies, 1993).

### *In situ* hybridization

Embryos were fixed in 4% PFA overnight at 4ºC. Embryos were dehydrated and rehydrated washing them with increasing and decreasing, respectively concentrations of methanol in PBT (25%, 50%, 75% and 100%). Bleaching was carried out by incubation in 6% H_2_O_2_ in PBT during one hour. Proteinase K (Sigma) digestion was performed at 10μg/ml with different incubation times depending on the stage. After permeabilization, embryos were washed with PBT during 5 minutes and fixed with glutaraldehyde 0.05% in 4%PFA during 20 minutes. Embryos were incubated in hybridization buffer (50% Formamide, 4x SSC pH 4.5, 1% SDS, 50μg/ml heparin (Sigma), 10μg/ml tRNA from baker’s yeast (Sigma), 1% w/v Blocking reagent (Sigma)) during 2 hours and hybridized with the probe overnight at 65ºC. Posthybridization washes were performed with 0.1% CHAPS w/v (Sigma), 2x SSC pH 5.5, followed by a second round of posthybridization washes with 0.1% CHAPS w/v, 0.2x SSC during 3 hours at 65ºC. Embryos were incubated overnight at 4ºC with 1:2000 anti-digoxigenin AP antibody (Roche) in 20% Goat serum, 1% Blocking reagent in TBST (5mM Tris-HCl pH 7.5, 15mM NaCl, 0.1% Triton X-100 (Sigma)). After several washes in TBST, embryos were washed with 125mM Tris-HCl pH 9.5, 125 mM NaCl, 62.5mM MgCl_2_, 0.5% Triton X-100 and stained with BMPurple (Roche) at room temperature until the signal was optimal. After the staining, embryos were washed with TBST, fixed in 4% PFA and stored at 4ºC. Occasionally, after *in situ* hybridization embryos were gelatin embedded and cryosectioned.

### Probe synthesis

RNA antisense probes were synthesized by transcription of linearized DNA from plasmids or from cDNA amplified with specific primers (S2 Table). Transcription was carried out with digoxigenin labelled nucleotides (Roche) and T7 RNA polymerase (Roche). Synthesized RNA was precipitated with 0.8M ammonium acetate in 75% ethanol or 0.1M LiCl in 75% ethanol and finally resuspended in 50% formamide-50% RNase free water.

### Victoria Blue staining

Embryos at E14.5 were eviscerated and fixed in 10% formaldehyde overnight and then washed in acid alcohol (3% HCl in 70% ethanol) several times. Embryos were stained during 3 hours with 0.5% w/v Victoria Blue (Sigma) in acid alcohol and after staining embryos were washed in acid alcohol until the embryos were white, then they were washed in 70% ethanol and 95% ethanol. Finally, embryos were clarified with increasing concentrations of Methyl salicylate in ethanol (30%, 50%) and stored in 100% Methyl salicylate.

### Alcian Blue and Alizarin Red staining

Embryos at E18.5 were eviscerated and the skin and soft tissues were removed as much as possible. Embryos were fixed overnight with 95% ethanol and after fixation were submerged in Alcian Blue solution (0.03% w/v Alcian Blue (Sigma), 80% ethanol, 20% glacial acetic acid) overnight. Alcian Blue solution was removed and several washes with 70% ethanol were made during the day; incubating the embryos in 95% ethanol overnight. Once the tissue becomes whiter, embryos were cleared with 1% KOH during 3-6 hours depending on the stage and the amount of soft tissue that the embryos have. Once cleared, Alizaren Red solution (0.005% Alizarin Red (Sigma), 1% w/v KOH) was added until the bones were stained. Another clarification step with 1% KOH could be done if necessary after staining with Alizarin Red solution, if not embryos were transferred to increasing concentrations of glycerol (20% and 50%) and finally placed in 100% glycerol for long term storage.

### Immunostaining and imaging

Embryos were fixed in 2% PFA, gelatin embedded and cryosectioned. Sections were permeabilized with 0’5% Triton X-100 in PBS for 20 minutes and blocking was performed with 20% goat serum in PBS for 1 hour. The primary antibodies used overnight at 4º were a rabbit anti-Aldh1a2 (ab96060) and an anti-Meisa, recognizing C-terminal short isoform of Meis1 and Meis2 (Mercader et al., 2005). Secondary antibodies were incubated during 45 minutes at room temperature. Secondary antibodies used were an Alexa-488 (1:500) for anti-Meis-a and an anti-HRP (1:200) for anti-Aldh1a2. After anti-HRP incubation, amplification with Tyr-FITC (1:100) during 3 minutes at room temperature was performed. Sections were incubated with DAPI and mounted in Vectashield or Dako fluorescent mounting media for acquisition. Images were acquired using a Nikon A1R confocal microscope using 405, 488 and 561nm wavelengths and Plan Apo 10x DIC L or Plan Apo VC 20x DIC N2 dry objectives.

### mRNA sequencing

Differential gene expression analysis was carried out between *Dll1^Cre^;Meis1^flox/flox^;Meis2^flox/flox^* and control embryos at E9. Four embryos were used for each condition and were staged by somite number, choosing the embryos with 20-24 somites. Total RNA was isolated using RNeasy Micro Kit (Qiagen) separating the anterior region containing the first 10-12 somites (the head was excluded) and the posterior region with the rest of the somites and the posterior embryonic bud. 20ng of total RNA were used to generate barcoded RNA-seq libraries using the NEBNext Ultra RNA Library preparation kit (New England Biolabs). The size and the concentration of the libraries were checked using the TapeStation 2200 DNA 1000 chip. Libraries were sequenced on a HiSeq2500 (Illumina) to generate 60 bases single reads. FastQ files for each sample were obtained using bcltofastQ software 2.20.

### RNA-seq data analysis

Sequencing reads were pre-processed by means of a pipeline that used FastQC (http://www.bioinformatics.babraham.ac.uk/projects/fastqc/), to asses read quality, and Cutadapt (Martin, 2011) to trim sequencing reads, eliminating Illumina adaptor remains, and to discard reads that were shorter than 30 bp. The resulting reads were mapped against the mouse transcriptome (GRCm38, release 91; dec2017 archive) and quantified using RSEM v1.2.20 (Li and Dewey, 2011). Data were then processed with a pipeline that used Bioconductor package Limma (Ritchie et al., 2015) for normalization and differential expression analysis, using a blocking strategy to consider gender and developmental stage (number of somites). Genes with at least 1 count per million in at least 4 samples (14,731 genes) were considered for further analysis. We considered as differentially expressed those genes with Benjamini-Hochberg adjusted p value <0.05. Fold change and log(ratio) values were calculated to represent gene expression differences between conditions.

## ACKNOWLEDGEMENTS

We thank members of the Torres group for stimulating discussions and suggestions. We thank members of the Genomics, Bioinformatics, Pluripotent Cell Technology, Transgenesis and Animal Facility CNIC units for excellent support. *Meis1* floxed mice were generated by Keith Humphries and kindly provided by Hesham Sadek. We thank Miguel Manzanares for providing mouse strains and Marian Ros, Aimee Zuniga, Paola Bovolenta, Robb Krumlauf and Jose Luis de la Pompa for kindly providing in situ probes.

## Author contributions

conceptualization: M.T. and A.C.L.; methodology: A.C.L., I.D. and V.C.; statistics: F.S.; investigation: A.C.L., V.C. and M.T.; writing (original draft): M.T. and A.C.L.; writing (review and editing): M.T., A.C.L.; supervision: M.T.; funding acquisition: M.T.

## Data and materials availability

All data needed to evaluate the conclusions in the paper are present in the paper and/or the Supplementary Materials except the RNA-seq data, which are available from GEO with reference number GSE146301. Materials are available from authors upon request.

---

**Figure S1.**
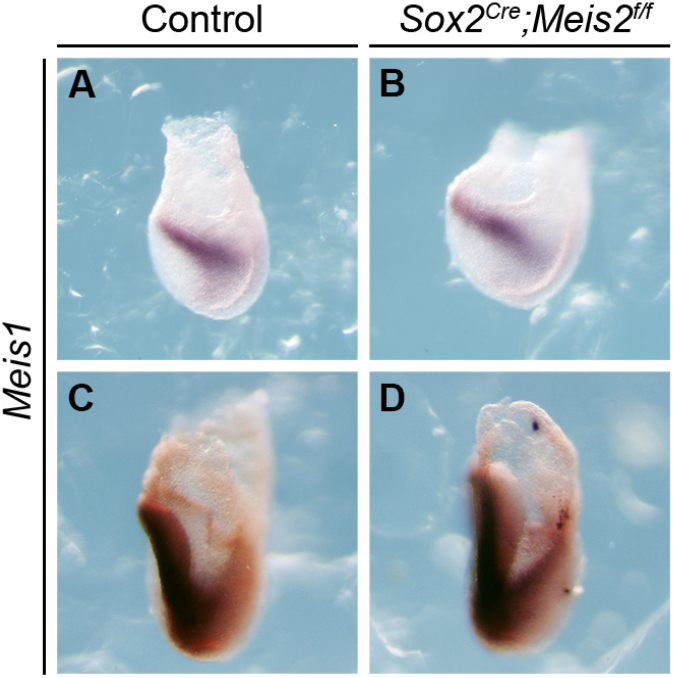
*Meis1* mRNA *in situ* hybridization in control and *Sox2^Cre^;Meis2^f/f^* embryos. (A and C) Control embryos at E7.5 and E8, respectively. (B and D) *Sox2^Cre^;Meis2^f/f^* embryos at E7.5 and E8, respectively.

**Figure S2.**
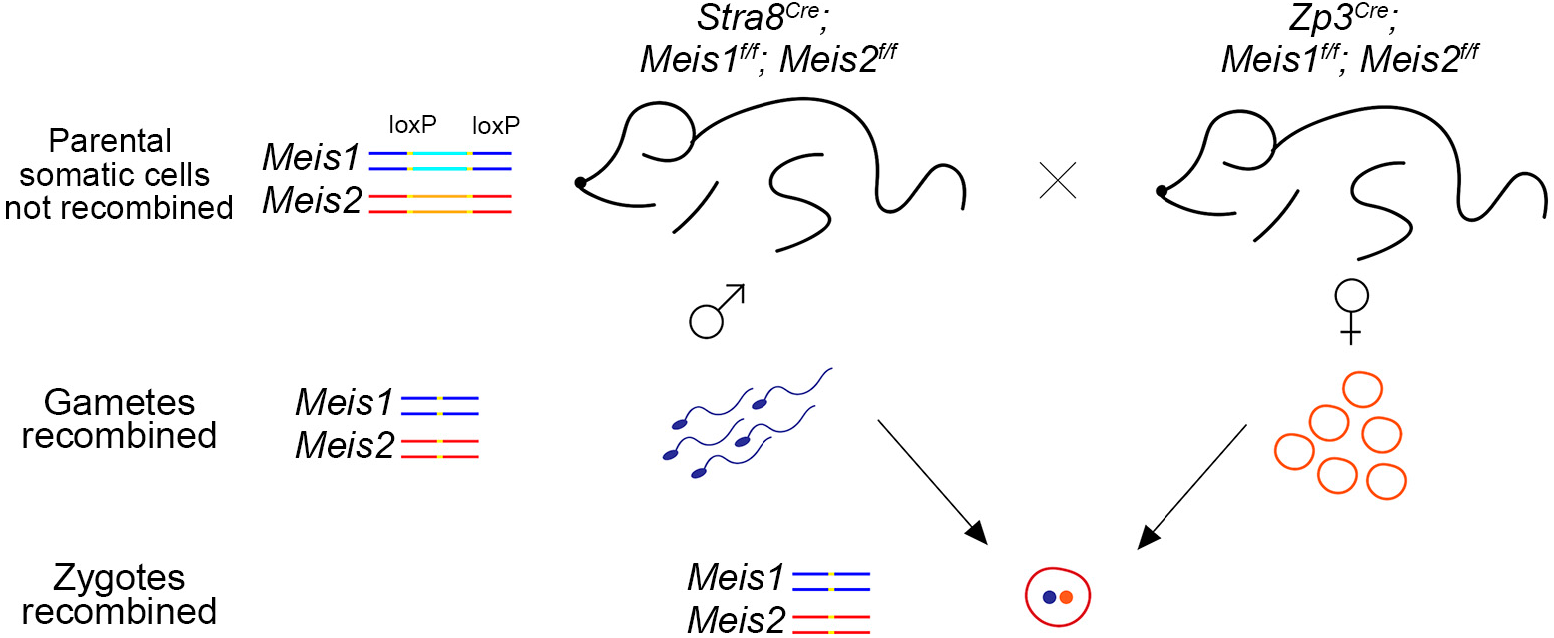
Schematic representation of crosses using biparental germ line Cre recombination to obtain complete zygotic elimination of *Meis1* and *Meis2*. *Meis1^f/f^;Meis2^f/f^* males and females respectively carrying *Stra8^Cre^* and *Zp3^Cre^* alleles only recombine floxed alleles in the germ line. Parental mice are viable while their progeny is double-knockout from the zygotic stage.

**Figure S3.**
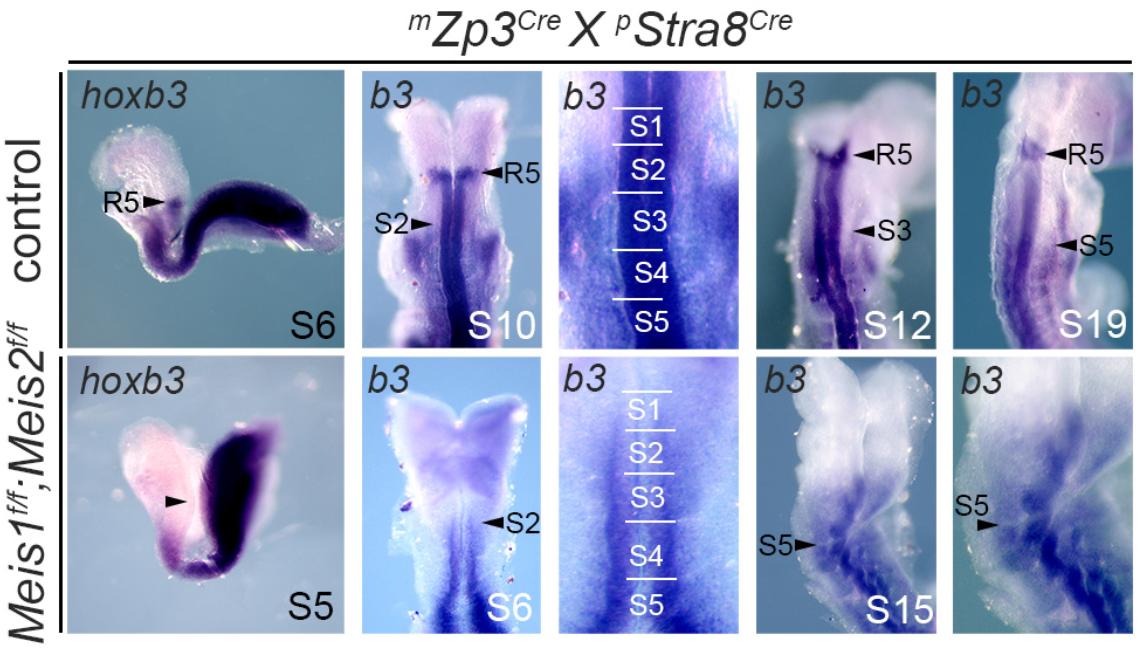
*hoxb3* gene mRNA expression patterns in *Meis* loss-of-function mutants. mRNA *in situ* hybridization of the indicated *Hox* genes in control and double-floxed *Meis1* and *Meis2* embryos derived from *Zp3^Cre^* mothers and *Stra8^Cre^* fathers.

**Figure S4.**
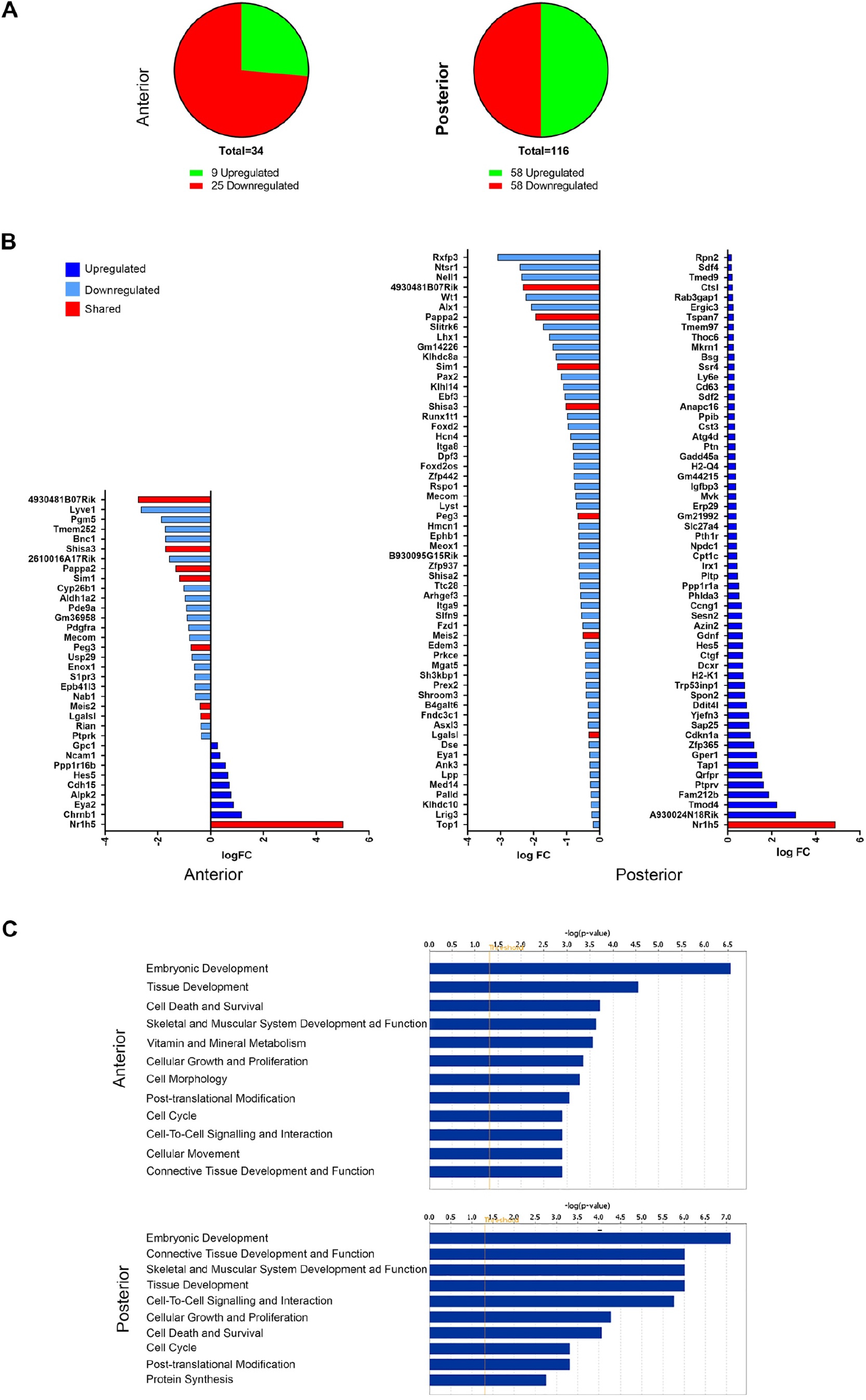
Comparative transcriptomic analysis of *Dll1^Cre^;Meis1^f/f^;Meis2^f/f^* and control embryos at E9. (A) Representation of the number of genes differentially expressed in both anterior and posterior samples (adjusted p-value ≤ 0.05). (B) Fold change representation (adjusted p-value ≤ 0.05) from anterior and posterior samples (upregulated and downregulated genes are colored in dark and light blue, respectively). Genes colored in red are differentially expressed in both, anterior and posterior. (C) Functions affected in *Dll1^Cre^;Meis1^f/f^;Meis2^f/f^* embryos from the Ingenuity Pathway analysis in anterior and posterior regions.

**Figure S5.**
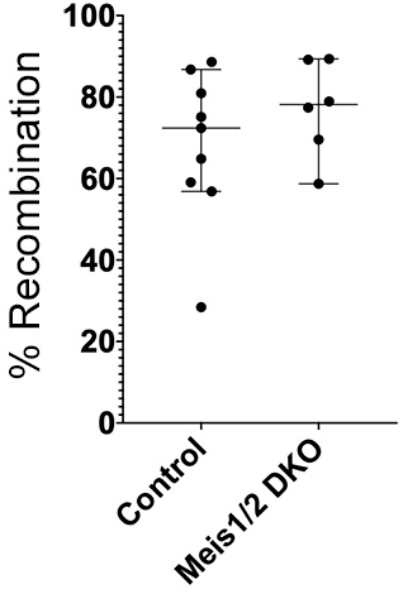
Frequency of recombination induced by *Dll1^Cre^* in the paraxial mesoderm. The graph shows the frequency of recombined cells measured in the 3 newly formed somites of E8.5-E10.5 *Dll1^Cre^;Rosa26R^Tomato^* embryos wild type for *Meis1* and *Meis2* (controls) or carrying the *Meis1^f^* and *Meis2^f^* alleles in homozygosity (Meis1/2 DKO). Graphs show individual measurements, the median and the 95% confidence interval.

**Figure S6.**
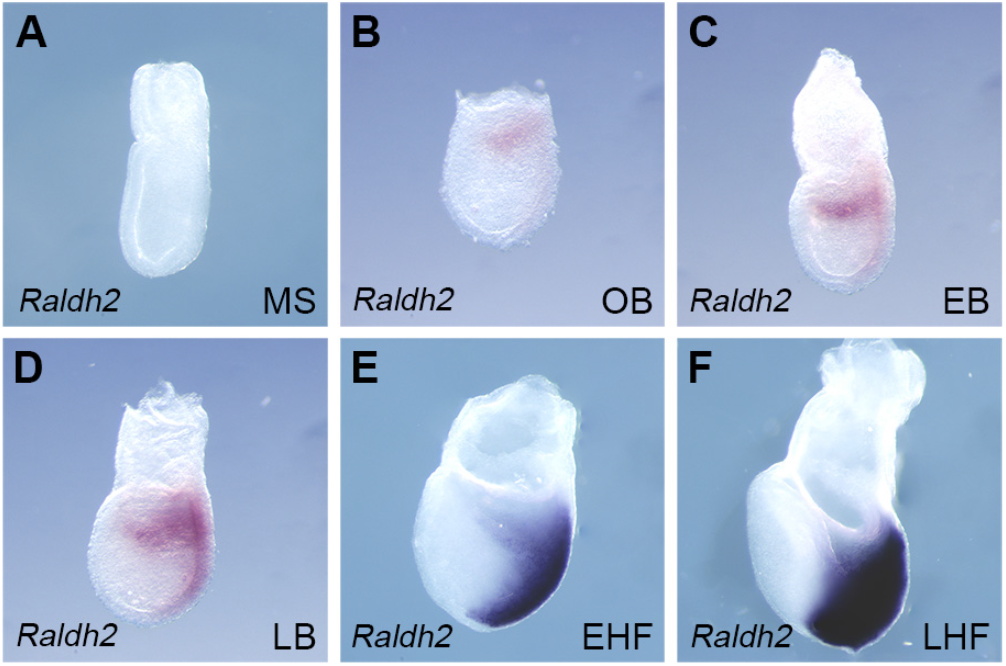
*Raldh2* expression pattern in early embryo. (A-F) Whole-mount mRNA *in situ* hybridization of *Raldh2* from E7 to E7.75. MS, mid-streak; OB, no allantoid bud; EB, early allantoid bud; LB, late allantoid bud; EHF, early headfold; LHF, late headfold. All images are oriented with the anterior to the left.

**Table S1A.** Scoring of skeletal defects in Conditional deletion of *Meis2* using different Cre drivers.

| **PHENOTYPES** | **Sox2Cre;M2-/-** | **Mesp1Cre;M2-/-** | **Dll1Cre;M2-/-** |
| --- | --- | --- | --- |
| **Abnormalities cervical vertebra** |  |  |  |
| C1 fused to E unilateral | 1 | 3 | 1 |
| C1 fused to E bilateral | 13 | 3 | 0 |
| C1 NA approaching E unilateral | 0 | 2 | 1 |
| C1 NA approaching E bilateral | 0 | 1 | 3 |
| C2 with C1-like morphology unilateral | 2 | 2 | 0 |
| C2 with C1-like morphology bilateral | 11 | 3 | 4 |
| C3 with C2-like morphology unilateral | 1 | 0 | 0 |
| C3 with C2-like morphology bilateral | 1 | 0 | 0 |
| Less than 7 cervical vertebra | 0 | 2 | 0 |
| Other abnormalities * | 10 | 4 | 3 |
| **Rib defects** |  |  |  |
| R1 short unilateral | 3 | 0 | 0 |
| R1 short bilateral | 5 | 0 | 3 |
| R1 short and fused R2 unilateral | 4 | 0 | 1 |
| R1 fused R2 unilateral | 0 | 0 | 0 |
| R1 fused R2 bilateral | 1 | 0 | 0 |
| R13 short | 6 | 1 | 2 |
| Other rib fusions/splits | 5 | 2 | 3 |
| Vestigial rib on L1 | 4 | 1 | 0 |
| sternal/floating ribs 6/7 unilateral | 1 | 0 | 2 |
| sternal/floating ribs R6/7 bilateral | 2 | 0 | 2 |
| **Sternum defects** | 5 | 1 | 0 |
| **Nº of embryos** | **14** | **9** | **4** |

The table shows the scoring of the observed skeletal defects. B, basioccipital; aaa, anterior arch of the atlas; C, cervical vertebra; NA, neural arches; E, exoccipital; Vb, vertebra; R, rib; L, lumbar vertebra. Number of studied embryos is indicated in parenthesis when it is different from the total.
\* Includes misshaping, split neural arches, extra-elements in the cervical region and mismatch in the posterior arch.

**Table S1B.**
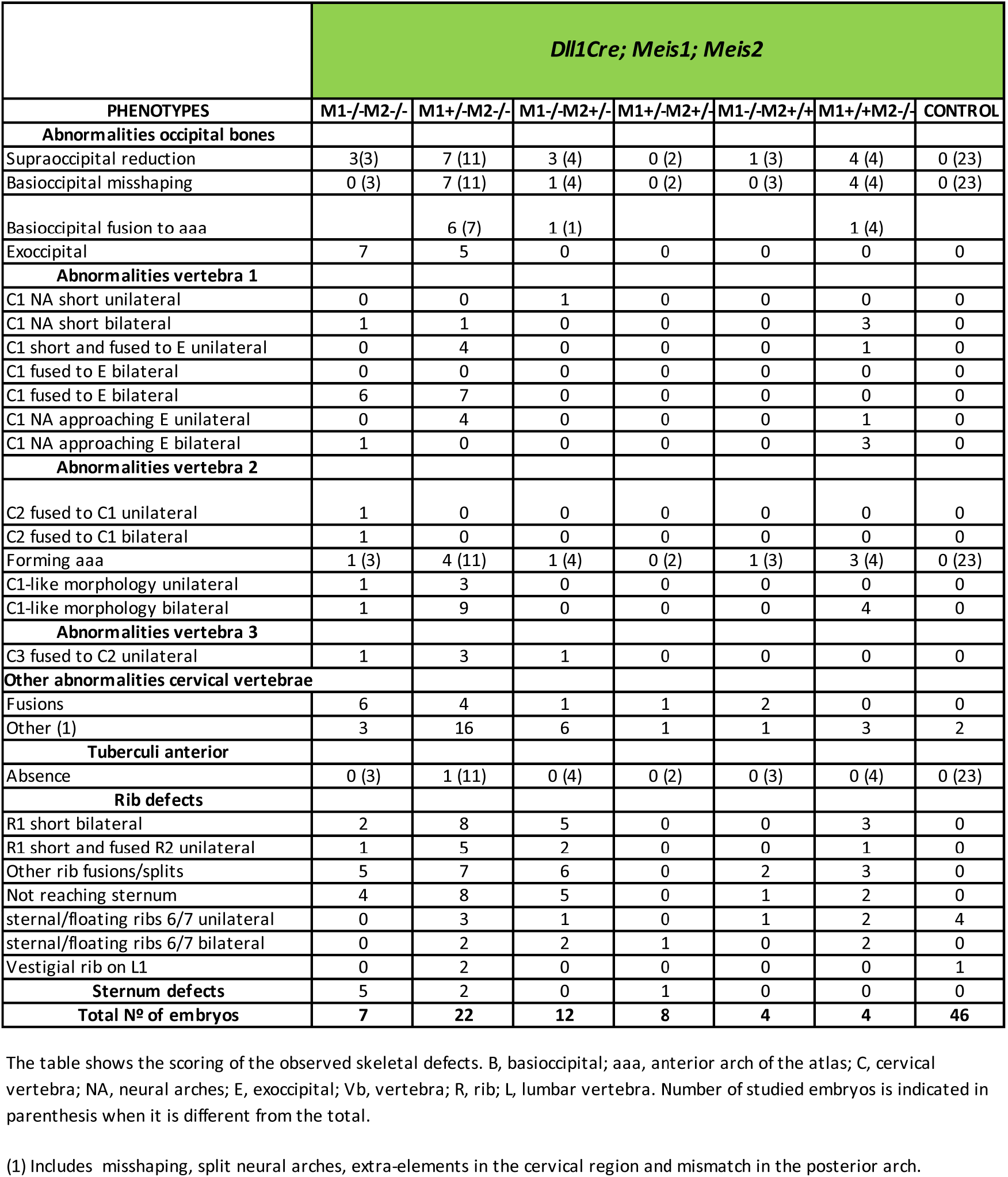
Scoring of skeletal defects in conditional deletion of *Meis1* and *Meis2* with *Dll1^Cre^*.

**Table S1C.** Statistical analysis of data in S1B.

|  | *Dll1*<sup>Cre</sup>; *Meis1*; *Meis2* |  |
| --- | --- | --- |
| PHENOTYPES | Control | M1-/M2+/- |
| **Any occipital defect** | 0(23) | 3(4)** |
| **Any cervical defect** | 2(46) | 4(12)* |
| **Any thoracic defect** | 4(46) | 10(12)* |

We used a Fisher test to compare proportions of given features between groups of fetuses of different groups
\* pvalue<0.05 ; \*\* pvalue<0.01

**Table S1D.** Scoring of skeletal defects in conditional deletion of *Raldh2* with *Dll1^Cre^*.

|  | *Dll1Cre; Raldh2* |  |  | *Dll1Cre; Raldh2;R26RMeis2* |
| --- | --- | --- | --- | --- |
| PHENOTYPES | -/- | +/- | +/+ | -/- |
| **Abnormalities occipital bones** |  |  |  |  |
| Supraoccipital reduction | 2 (25) | 0 (8) | 4 (33) | 0(6) |
| Basioccipital misshaping | 14 (43) | 0 (8) | 5 (39) | 0(6) |
| Basioccipital fusion to aaa | 7 (14) |  | 4 (5) | 0(6) |
| Exoccipital misshaping | 3 | 0 | 0 | 0(6) |
| **Abnormalities vertebra 1** |  |  |  |  |
| C1 NA short unilateral | 5 | 0 | 0 | 0(6) |
| C1 NA short bilateral | 2 | 0 | 0 | 0(6) |
| C1 short and fused to E unilateral | 1 | 0 | 0 | 0(6) |
| C1 fused to E unilateral | 2 | 0 | 0 | 0(6) |
| C1 fused to E bilateral | 3 | 0 | 0 | 0(6) |
| C1 NA approaching E unilateral | 2 | 0 | 0 | 0(6) |
| **Abnormalities vertebra 2** |  |  |  |  |
| C2 fused to C1 unilateral | 8 | 0 | 2 | 0(6) |
| C2 fused to C1 bilateral | 1 | 0 | 0 | 0(6) |
| Forming aaa | 14 (41) | 0 (8) | 0 (38) | 0(6) |
| C1-like morphology unilateral | 2 | 0 | 0 | 0(6) |
| C1-like morphology bilateral | 6 | 0 | 0 | 0(6) |
| **Abnormalities vertebra 3** |  |  |  |  |
| C3 fused to C2 unilateral | 10 (41) | 0 (8) | 1 (38) | 0(6) |
| C3 fused to C2 bilateral | 1 (41) | 0 (8) | 0 (38) | 0(6) |
| **Abnormalities cervical vertebra** |  |  |  |  |
| Fusions | 10 | 0 | 0 | 0(6) |
| Other (1) | 12 | 1 | 2 | 0(6) |
| **Tuberculi anterior** |  |  |  |  |
| Absence | 1 (21) | 0 (8) | 0 (25) | 1(6) unilateral |
| Relocated to Vb7 unilateral | 4 (21) | 0 (8) | 2 (25) | 0(6) |
| Relocated to Vb7 bilateral | 1 (21) | 0 (8) | 0 (25) | 0(6) |
| **Rib defects** |  |  |  |  |
| R1 short unilateral | 2 | 0 | 0 | 0(6) |
| R1 short and fused R2 unilateral | 2 | 0 | 0 | 0(6) |
| R1 fused R2 bilateral | 5 | 0 | 1 | 0(6) |
| Other rib fusions/splits | 7 | 0 | 0 | 0(6) |
| Not reaching sternum | 4 | 0 | 0 | 0(6) |
| sternal/floating ribs 8/5 unilateral | 7 | 0 | 5 | 2(6) |
| sternal/floating ribs 8/5 bilateral | 8 | 1 | 9 | 1(6) |
| sternal/floating ribs 6/7 unilateral | 1 | 0 | 0 | 0(6) |
| vestigial rib on L1 | 5 | 0 | 2 | 1(6) unilateral |
| **Sternum defects** | 1 | 0 | 0 | 0(6) |
| **N° of embryos** | **49** | **15** | **42** | **6** |

The table shows the scoring of the observed skeletal defects. B, basioccipital; aaa, anterior arch of the atlas; C, cervical vertebra; NA, neural arches; E, exoccipital; Vb, vertebra; R, rib; L, lumbar vertebra. Number of studied embryos is indicated in parenthesis when it is different from the total.
(1) Includes misshaping, split neural arches, extra-elements in the cervical region and mismatch in the posterior arch.

**Table S1E.** Statistical analysis of data in S1D.

|  | *Dll1Cre; Raldh2* |  | *Dll1Cre; Raldh2;R26RMeis2* |
| --- | --- | --- | --- |
| PHENOTYPES | +/- OR +/+ | -/- | -/- |
| **Any cervical defect** | 6(58) | 33(49)*** | 0(6)** |
| **Cervical transformations** | 3(58) | 24(49)*** | 0(6)* |
| **Any thoracic defect** | 7(58) | 33(49)*** | 0(6)** |

We used a Fisher test to compare proportions of given features between groups of fetuses
\* pvalue<0.05 ; \*\* pvalue<0.01 ; \*\*\*pvalue<0.001

**Table S2.**
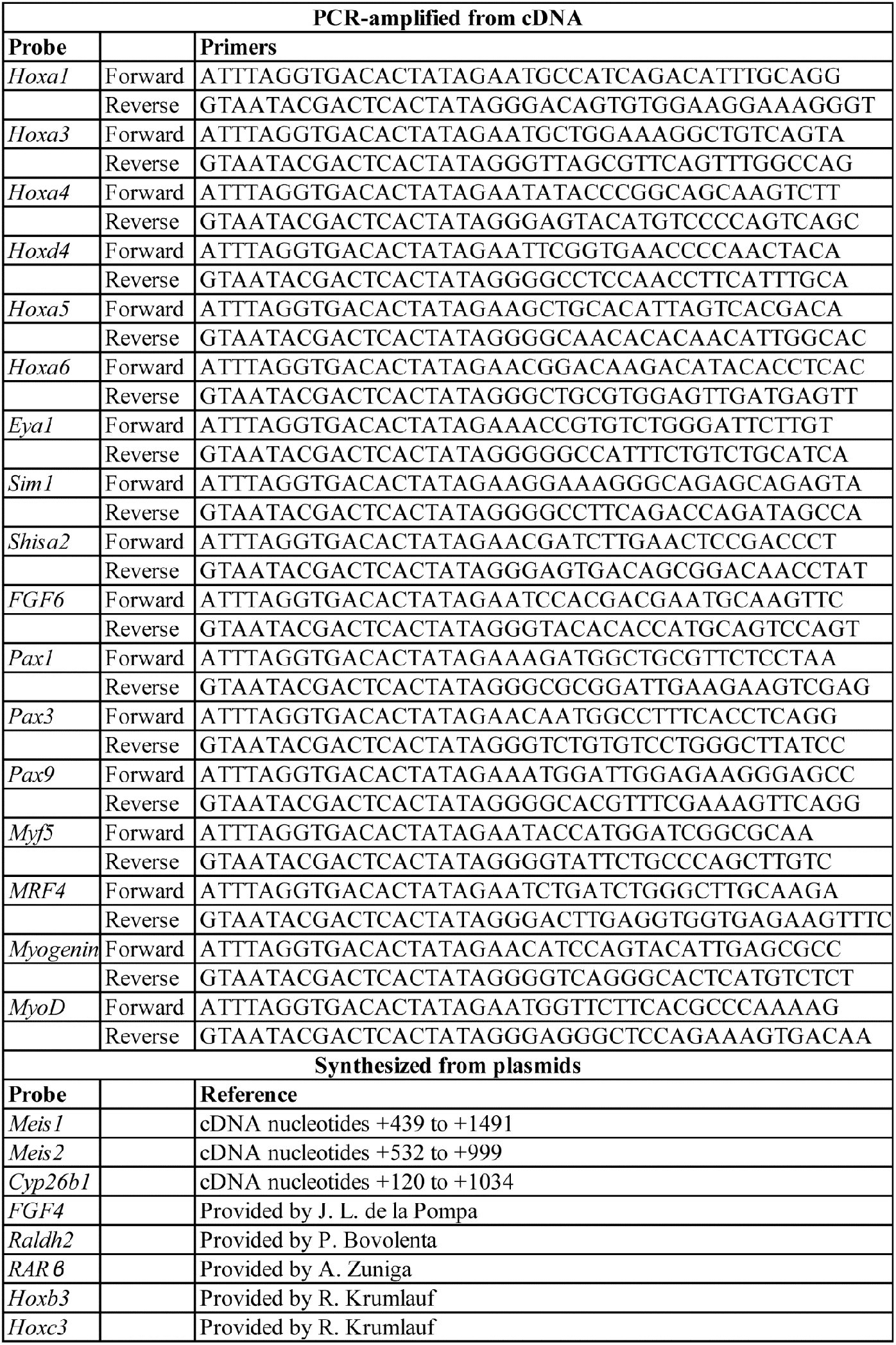
Probes used for whole-mount in situ hybridization.

